# Stromal transdifferentiation drives lymph node lipomatosis and induces extensive vascular remodeling

**DOI:** 10.1101/2022.06.30.498248

**Authors:** Tove Bekkhus, Anna Olofsson, Ying Sun, Peetra Magnusson, Maria H. Ulvmar

## Abstract

Lymph node (LN) lipomatosis is a common, but rarely discussed phenomenon, associated with aging, involving a gradual exchange of the LN parenchyma into adipose tissue. The mechanisms behind these changes and the effects on the LN have been unknown. We show that LN lipomatosis starts in the medullary regions of the human LN and link the initiation of lipomatosis to transdifferentiation of LN medullary fibroblasts into adipocytes. The latter is associated with a downregulation of lymphotoxin beta expression. We also show that, medullary fibroblasts, in contrast to the reticular cells in the T-cell zone, display an inherent higher sensitivity for adipogenesis. Progression of lipomatosis leads to a gradual loss of the medullary lymphatic network, but at later stages, collecting-like lymphatic vessels, are found inside the adipose tissue. The stromal dysregulation includes a dramatic remodeling and dilation of the high endothelial venules associated with reduced density of naïve T-cells. Abnormal clustering of plasma cells is also observed. Thus, LN lipomatosis causes widespread stromal dysfunction with consequences for the immune contexture of the human LN. Our data warrant an increased awareness of LN lipomatosis as a factor contributing to decreased immune functions in the elderly and in disease.

**Graphical abstract:** In lymph nodes (LNs) of young patients there is a normal lymph flow, normal and functioning high endothelial venules (HEVs) with a high density of surrounding naïve T-cells. With aging lymphotoxin beta (LTB) is downregulated in the medulla of the LN and the fibroblasts of the medulla, namely the medullary reticular cells (MedRCs), transdifferentiate into adipocytes inducing LN lipomatosis. LN lipomatosis leads to loss of lymphoid tissue, medullary sinuses and can be predicted to result in a shortcut of the lymph flow based on the presence of collecting-like vessels in the adipose tissue in late stage lipomatosis. Lipomatosis also induce extensive vascular remodeling with loss of medullary lymphatic vessels and dysfunctional, highly dilated HEVs with lower density of naïve T-cells and trapped plasma cells.

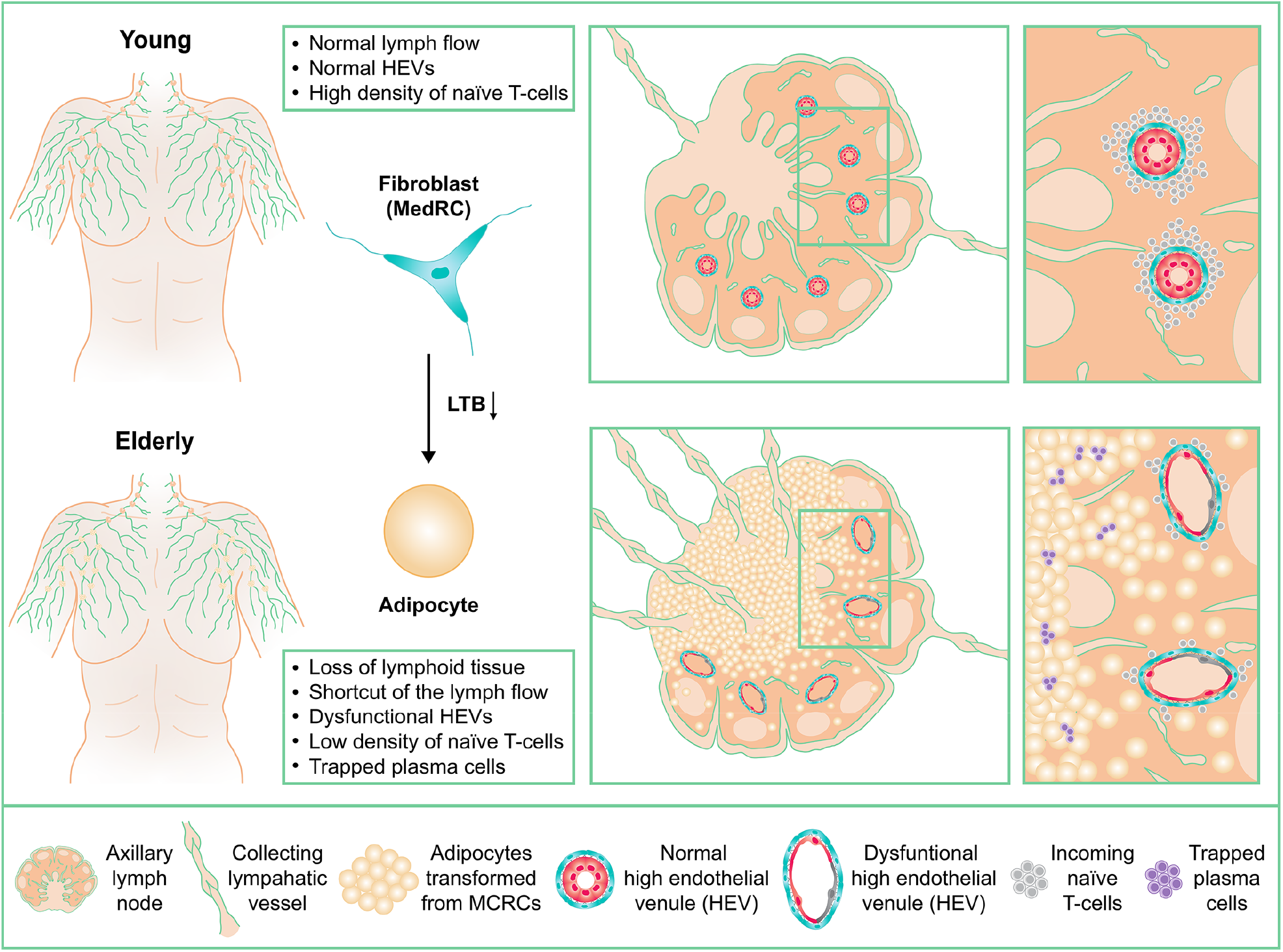

## Introduction

With aging, the function of our immune system decline and it is well established that for example vaccinations do not work as well in elderly as in the younger [1]. The underlying reasons for impaired immune responses in the elderly are complex and most likely multifactorial. One group of contributing factors can be linked to environmental changes in lymphoid tissues [2]. This includes involution of the thymus [3,4] and a decline in bone marrow (BM) hematopoietic stem cells [2]. In both these cases, the healthy tissue is eventually replaced by adipocytes [2-4]. A related but less well-known and studied process is lymph node (LN) lipomatosis [5,6]. Similar to the age related accumulation of adipocytes in thymus and BM [2,4], LN lipomatosis is a process where the normal parenchyma is replaced by adipocytes. Analysis of multiple LNs across the human body support that lipomatosis increase progressively with age [5-7].

The LNs are crucial organs for the induction of protective adaptive immune responses and immunological memory in vaccination, infection and cancer [8,9]. They are characterized by highly structured immune cell compartments allowing blood derived naïve B- and T-cells to become activated by lymph derived antigens and antigen presenting cells (APCs) [8]. The structure and function of the LN depends on its highly specialized stromal cell environment with LN specific subsets of fibroblasts [10] and a highly specialized blood and lymphatic vasculature [8,11-13]. The high endothelial venules (HEVs), which constitutively express a unique pattern of L-selectin ligands called peripheral node addressins (PNAd) [14], are essential for lymphocyte recruitment into the LN. The LN sinusoidal lymphatic sinuses allow transport of tissue antigens and APCs into the LN but also provide the exit route out of the LN for activated and recirculating lymphocytes [8,15,16]. We and others have also recently shown that the LN lymphatic vasculature display impressive heterogeneity and niche specific functions [11,12], reflecting the need to coordinate complex immune trafficking and responses.

Although LN lipomatosis is common pathological change seen in the elderly [5], the mechanisms driving these changes and the potential effects on the LN microenvironment have up until now not been studied. As LNs are always surrounded by adipose tissue [17], one possible mechanism that have been proposed is the infiltration of surrounding adipocytes into the LN [7]. Our data do not support this notion. Instead, by careful analysis of LNs with different degree of lipomatosis we can show that it starts from deeper parts of the medullary parenchyma, and demonstrate the presence of cells that display a transitional phenotype with both fibroblast and adipocyte lineage marker expression. This suggests that LN lipomatosis is driven by transdifferentiation of medullary fibroblasts into adipocytes. We show that these changes are associated with downregulation of lymphotoxin beta (*LTB*) in affected areas, a factor known to counteract adipocytic differentiation of stromal cell precursors in early development of the LN [18]. We also provide evidence that supports that medullary reticular fibroblast (MedRCs) are inherently more sensitive to transdifferentiate into adipocytes, than T-cell zone recticular cells (TRCs), providing an additional explanation to the observed initiation of the pathology in the medulla of the LN. Finally, we demonstrate that LN lipomatosis causes loss of the medullary stroma and extensive vascular remodeling of the HEVs and the lymphatic vasculature, changing the immune contexture of the human LN.

## Results

### Lipomatosis is frequent in lymph nodes from elderly patients

Lymph node (LN) lipomatosis is common in humans above 40 years of age and is progressively more frequent with aging [5,6]. To analyze the pathology of LN lipomatosis, we used a collection of axillary LNs without metastasis from breast cancer patients [19], including patients with non-invasive ductal carcinoma in situ (DCIS) and invasive ductal carcinoma (IDC)), and a subset of pancreatic LNs from cancer free organ donors (ODs). LNs were defined as having lipomatosis based on the presence of parenchymal adipocytes inside of the LN capsule (Figure 1A and 1B). A majority of the patients displayed LN lipomatosis (Figure 1B). Figure 1C illustrates an inter- and intra-patient variation in our analyzed patient cohort, where some LNs had no lipomatosis and others had lost all normal parenchyma.

**Figure 1:**
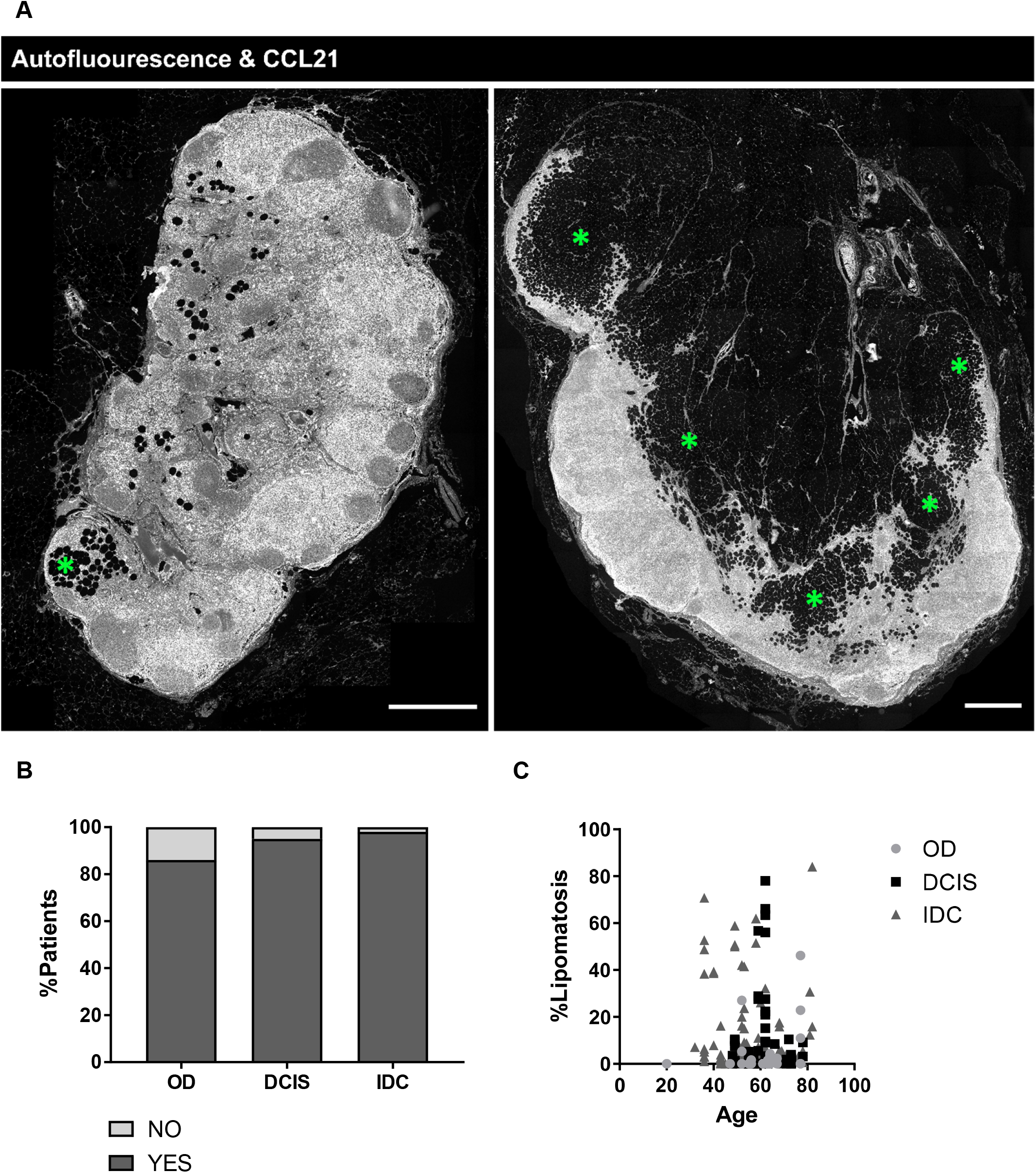
Lipomatosis is frequent in lymph nodes from elderly patients. **(A)** Immunofluorescence staining of axillary lymph nodes (LNs) with low (left, 5.4%) and high (right, 62.0%) lipomatosis burden. Staining of the chemokine CCL21 in the green channel allowing structural visualization through autofluorescence. Green asterisks marks examples of adipocytes. Scale bar: 1mm. **(B)** Frequency of the presence of lipomatosis in organ donors (ODs, n=14 donors), patients with ductal carcinoma in situ (DCIS, n=22 patients) and invasive ductal carcinoma (IDC, n=44 patients). **(C)** Plot of lipomatosis burden (% lipomatosis area/LN area) and age of the patients including OD (n=12 donors, n=57 LNs), DCIS (n=20 patients, n=59 LNs), IDC (n=37 patients, n=93 LNs). Every dot represents one LN (n=209). The age range was similar between the groups: i.e. OD: 57±14, DCIS: 61±12 and IDC: 57±12.

### Lipomatosis starts within the medullary compartment and is driven by adipogenic transdifferentiation of medullary fibroblasts

It is not established if development of lipomatosis is more frequent within specific areas of the LN. With this in mind we analyzed the localization of the lipomatosis based on the basic structure of the LN; cortex (B-cell area), paracortex (T-cell area), and medulla (central lymphatic sinus area) [20]. Lipomatosis was most frequently found within the medullary compartment and was only found in the paracortex and/or cortex if the lipomatosis burden was large enough to also reach these parts of the LN (Figure 2A-B, also see Figure 1A; right panel). In LNs with a low or intermediate degree of lipomatosis, it was exclusive to the medullary region and could be seen randomly across the medulla (Figure 2B, also see Figure 1A; left panel). This is supports neo-formation of adipocytes within the medulla rather than invasion of the adipocytes surrounding the LN. The latter would be expected to first affect peripheral areas including the cortex rather than deep areas of the LN parenchyma.

**Figure 2:**
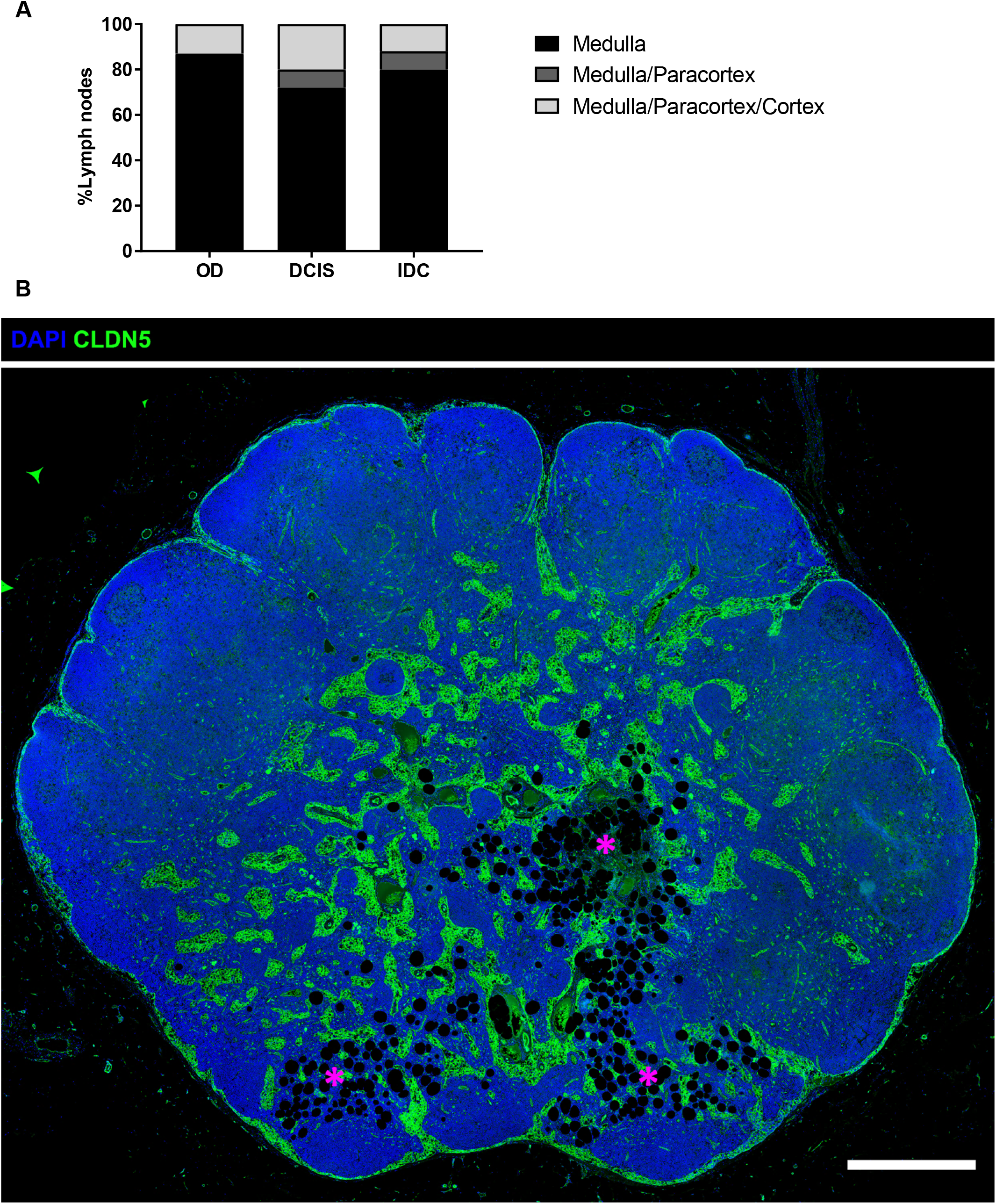
Lipomatosis starts within the medullary compartment. **(A)** Localization frequency of adipocytes in lymph nodes (LNs) from organ donors (ODs, n=8 donors, n=15 LNs), patients with ductal carcinoma in situ (DCIS, n=21 patients, n=54 LNs) and invasive ductal carcinoma (IDC, n=44 patients, n=107 LNs). **(B)** Immunofluorescence staining of an axillary LN with intermediate lipomatosis localized to the medulla marked by asterisks in magenta. LN stained for the vascular tight junction protein Claudin-5 (CLDN5, green) marking the medullary lymphatic sinuses together with DAPI (nuclei). Scale bar: 750μm. Representative image is shown.

Experimental data in mice support a sensitive balance in the differentiation of LN mesenchymal cells into fibroblast lineages or adipocytes that depend on lymphotoxin beta receptor (LTBR) signaling [18]. To evaluate if differentiation of medullary fibroblasts into adipocytes could be one contributing factor in driving lipomatosis, we stained for Perilipin, which is an marker of adipocyte differentiation [21], together with alpha smooth muscle actin (αSMA), a general marker for fibroblasts that is not expressed by adipocytes, and Claudin-5 (CLDN5), a tight junction protein expressed by several types of endothelial cells, including the lymphatic endothelial cells (LECs) forming the medullary sinuses, but not by fibroblasts or adipocytes (Figure 3A-C) [11]. Sometimes the adipocytes appeared to transverse into the sinuses between the fibroblasts that are lining the medullary sinuses (arrowheads in Figure 3A). Perilipin and αSMA could be observed to be co-expressed in individual cells in all the analyzed LNs (Figure 3B, Supplementary Figure 1). CLDN5 or LYVE-1 was in contrast never found to be co-expressed with Perilipin (Figure 3C). Taken together, the dual expression of αSMA and Perilipin in the same cells is suggestive of the presence of medullary fibroblast in transition into adipocytes.

**Figure 3:**
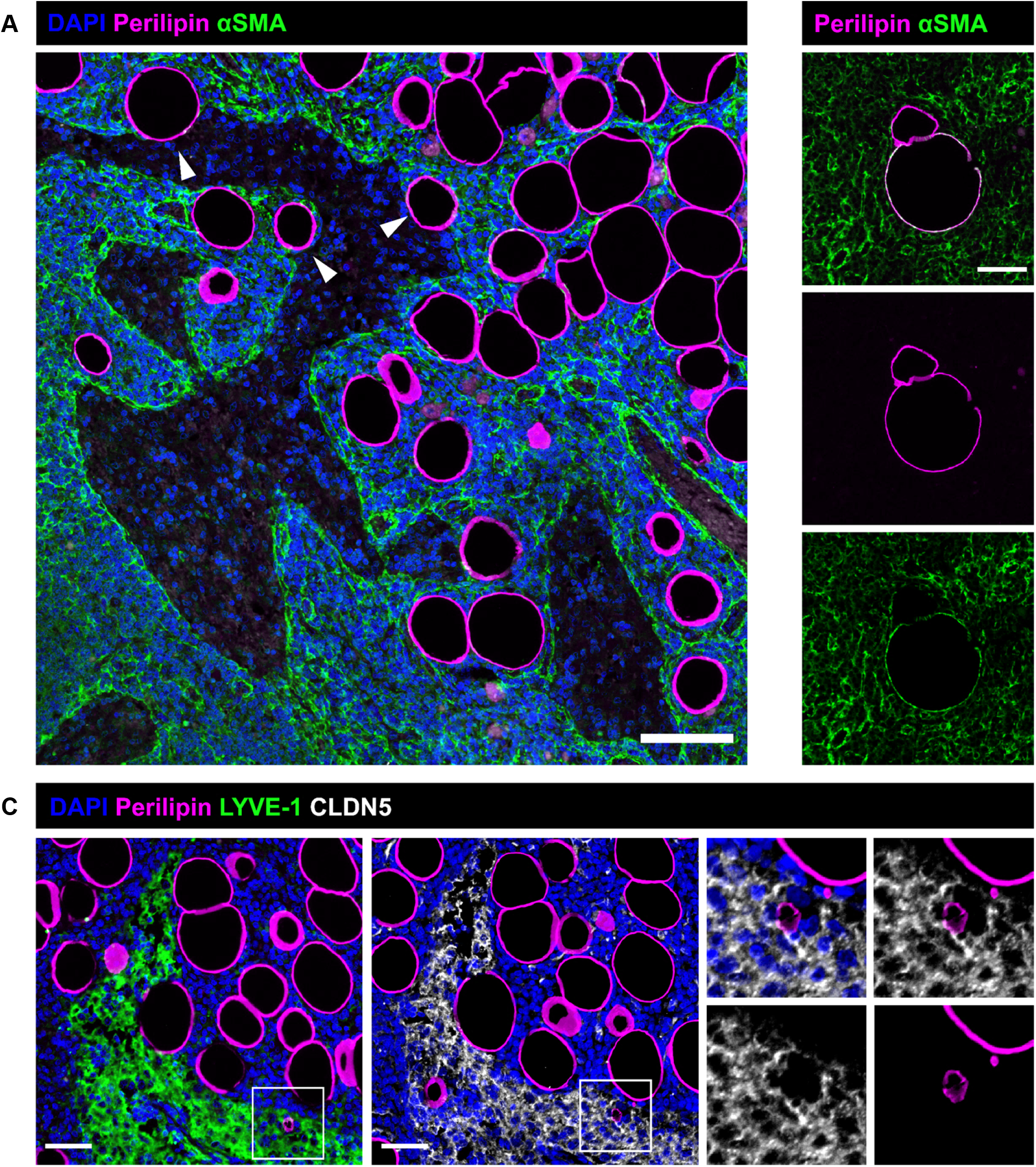
Lipomatosis is driven by adipogenic transdifferentiation of medullary fibroblasts. Immunofluorescence staining of lymph nodes (LNs) with lipomatosis. **(A)** Presence of adipocytes (Perilipin, magenta) and fibroblasts (alpha smooth muscle actin; αSMA, green) in the medullary cords. Arrowheads point out adipocytes present in the layer of medullary lymphatic sinus lining fibroblasts. Scale bar: 100μm. Pictures representative for 10 analyzed LNs. **(B)** Co-staining of αSMA (green) and Perilipin (magenta) in medullary fibroblasts. Scale bar: 50μm. Pictures representative for 10 analyzed LNs. **(C)** Presence of adipocytes (Perilipin, magenta) within the medullary sinus (LYVE-1, green and Claudin-5; CLDN5, grey) that do not co-stain with neither LYVE-1 (green) nor CLDN5 (grey). Pictures representative for 6 analyzed LNs. Scale bar: 50μm.

### Cell intrinsic sensitivity to adipogenesis of CD34^+^ SCs and MedRCs

The transdifferentiation of LN fibroblasts into adipocytes seen specifically in the medulla may reflect that the medullary area contain specific phenotypes of fibroblasts more prone to convert into an adipogenic phenotype and/or that the LN environment in this area change in aging, driving adipogenic transdifferentiation of fibroblasts in this specific region, or a combination of both. A niche specific heterogeneity is known to characterize the LN fibroblasts [10,22]. The main subsets include the follicular dendritic cells (FDCs), organizing the cortical B-cell zone of the LN and the T-cell reticular cells (TRCs), organizing the paracortex. Two subsets of medullary fibroblast, here referred to as medullary reticular cells (MedRCs) and one subset of CD34^+^ fibroblasts, associated with the capsule and adventitia of large blood vessels, have also been identified [10]. While most current mapping and molecular characterization of the fibroblast subsets in LNs are derived from studies of mouse [10,23], recent single cell analyses of human LN fibroblasts indicates that major functions and subsets are conserved [22,24]. By staining human LNs (with low lipomatosis) we confirm on the protein level that this include a conserved higher CCL21 expression in TRCs compared to MedRCs (Supplementary Figure 2) and that CD34^+^ fibroblasts, similar to mouse [10], mainly is associated with blood vessels, the capsule and the adventitia of large blood vessels (Supplementary Figure 3).

To evaluate the potential difference between LN fibroblast subsets in sensitivity to adipogenesis we took use of the difference in expression of the marker BP3 and the marker CD34 in mouse LN fibroblasts [10,23]. While TRCs and FDCs express high BP3, it is low in MedRCs [23]. By adding the marker CD34, it is possible to sort three different subsets of podoplanin (PDPN) positive SCs from the mouse LN (Supplementary Figure 4). These include BP3^pos^ CD34^neg^ PDPN^pos^ cells, which will be dominated by TRCs but that may also contain the less abundant populations FDCs and marginal reticular cells (MRCs), (from here on therefore referred as BP3^+^ RCs); the BP3^neg^ CD34^neg^ PDPN^pos^ cells which correspond to MedRCs; and last CD34^pos^ BP3^neg^ PDPN^pos^ cells, here referred to as CD34^+^ SCs. Under adipogenic culture conditions, CD34^+^ SCs displayed the highest conversion into adipocytes, followed by MedRCs while the BP3^+^ RCs displayed minimal conversion (Figure 4A-B). This supports that MedRCs and CD34^+^ SCs share an ability to convert into adipocytes while BP3^+^ RCs are inherently resistant under these conditions.

**Figure 4.**
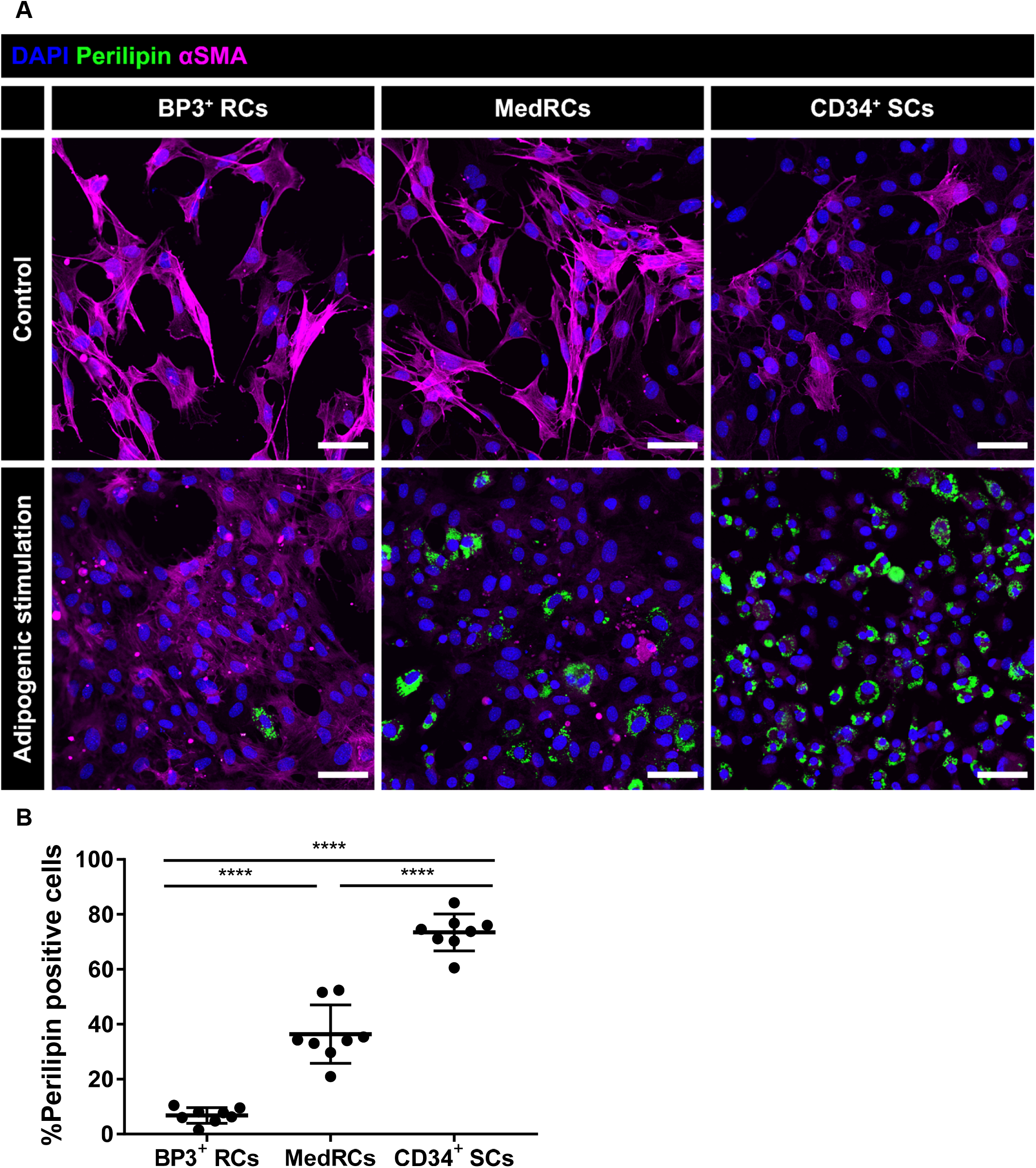
Cell intrinsic sensitivity to adipogenesis of CD34^+^ SCs and MedRCs compared to other lymph node reticular cell subsets. **(A)** Immunofluorescent staining of cultured fibroblast subsets; BP3^+^ reticular cells (RCs), medullary reticular cells (MedRCs) and CD34^+^ SCs, including both control (in non-adipogenic culture for 3 days) and adipogenesis stimulated cells (in non-adipogenic culture for 3 days and in adipogenic stimulation culture for 3 days). DAPI (blue), Perilipin (green) and alpha smooth muscle actin (αSMA, magenta). Scale bar: 50μm. **(B)** Quantification of the percentage of Perilipin positive cells after 3 days with adipogenic stimulation. Eight frames (mean: 118 cells/frame) per subset were analyzed and the experiment was repeated once. Lines represent mean and standard deviation. One-way ANOVA and Holm-Sidak’s multiple comparisons test was used for statistical analysis. ****p<0.0001.

To further understand these differences we used the published single cell analysis by Rodda *et al*. [10], to re-analyze the mouse LN fibroblasts and focused on differential expressed genes in the NF-kappa B signaling pathway, and the Hippo pathway, both implicated in controlling cell fate decisions between LN fibroblast or adipocytes [18,25] (Figure. EV6A-C). For analysis, cell types were stratified based on expression of bone marrow stromal cell molecule *Bst1*, also known as *Bp3*. TRC subsets, MRCs and FDCs were defined as *Bst1/Bp3* high population. Cell types of “Nr4a1^+^ SC”, “Inmt^+^ SC” and “CD34^+^ SC” were defined as *Bst1/Bp3* low populations. All these subsets express similar levels of *Ltbr* (Supplementary Figure 4). However, the analysis shows that *Bst1/Bp3*^+^ subsets; TRCs, MRCs and FDCs are enriched for genes associated with the NF-kappa B signaling pathway (including e.g. *Relb* and *NFkb2*), downstream of LTBR, while Nr4a1^+^ SCs and Inmt^+^ SCs (both MedRCs, that have been described to be located close to the medullary cords [10]) together with CD34^+^ SCs have low expression (Supplementary Figure 5). It can also be noted that TRC subsets, MRCs and FDCs display higher expression of the LTBR ligand *lymphotoxin beta (Ltb*) compared to MedRCs and CD34^+^ SCs (Supplementary Figure 5). Similar results were observed for genes associated with the Hippo pathway (Supplementary Figure 5). These data support cell intrinsic differences in both these pathways, associated with resistance to adipogenic transformation of LN fibroblasts.

**Figure 5:**
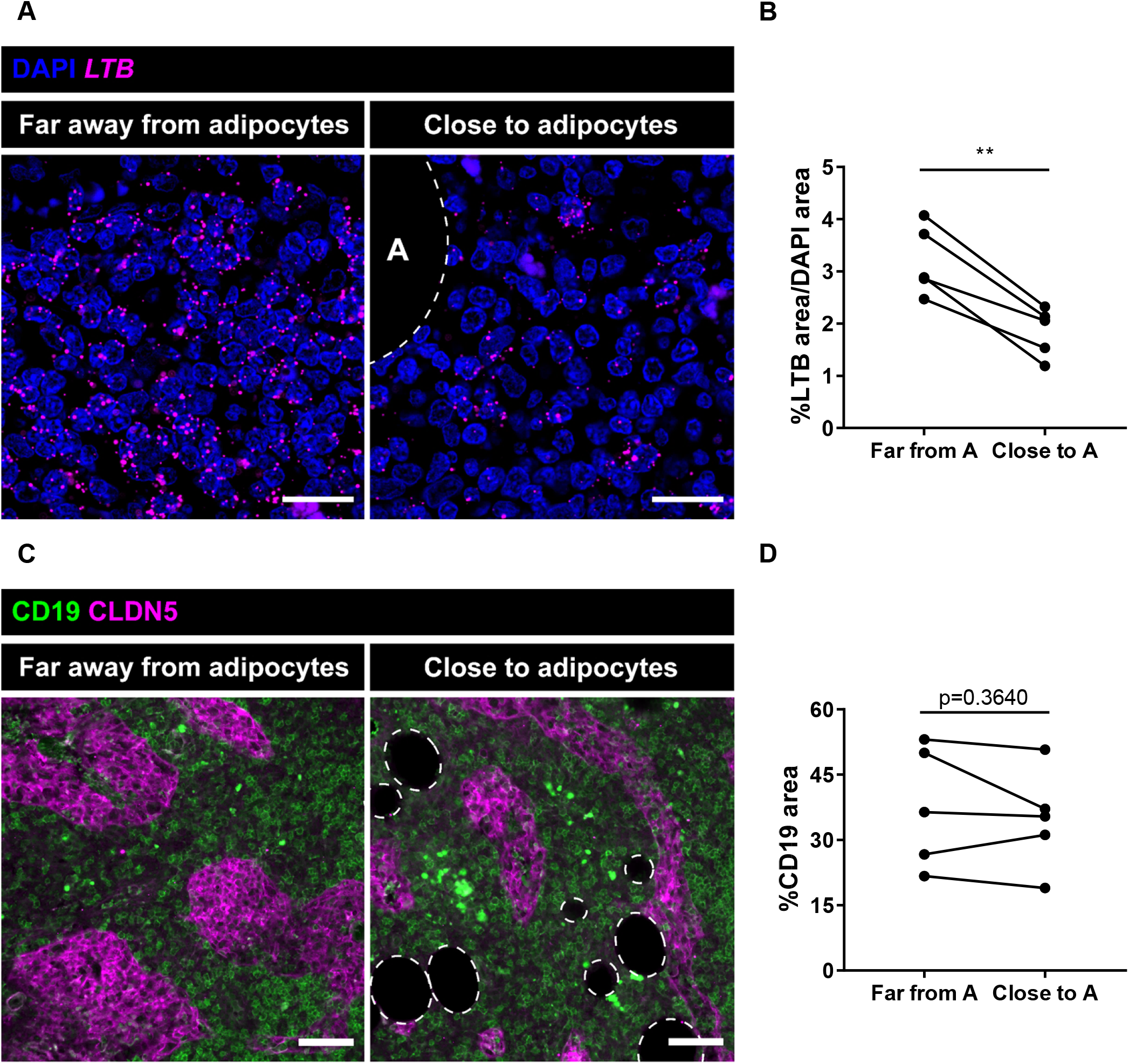
Lymph node lipomatosis is associated with downregulation of lymphotoxin beta in affected areas. **(A)** RNAscope of lymphotoxin beta (LTB) mRNA in the medulla far away from adipocytes (left) and close to adipocytes (right). Pictures taken from the same lymph node (LN). Dotted line and A visualizes the presence of adipocytes. Scale bar: 20μm. **(B)** Quantification of the signal of LTB mRNA as a percentage of LTB mRNA area out of DAPI area in parts of the medulla far away from adipocytes vs. close to adipocytes within the same LNs (n=5 LNs). Dots represent the mean value of three frames per LN. Paired t-test was used for statistical analysis. **p<0.01. **(C)** Immunofluorescent staining of CD19 (medullary B-cells, green) and Claudin-5 (CLDN5) (medullary lymphatic sinuses, magenta) in parts of the medulla far away from adipocytes and close to adipocytes within the same LN. Dotted lines visualize adipocytes. Scale bar: 75μm. **(D)** Quantification of the density of medullary B-cells as the percentage of the medullary area not covered by adipocytes or lymphatic sinuses within the same LN (n=5 LNs). Dots represent the mean value of three frames per LN. Paired t-test was used for statistical analysis. No significant difference was detected between the groups.

### Lymph node lipomatosis is associated with downregulation of LTB in affected areas

LTB can counteract adipogenesis of LN stromal cells. We therefore next analyzed *LTB* expression in LNs with early stage, low to intermediate lipomatosis, comparing areas affected or not affected by lipomatosis within the same LNs. Lower levels of *LTB* was detected in medullary areas associated with lipomatosis compared to the areas without adipocytes (Figure 5A-B). B-cells are known to be a main source of LTB in the LN [26]. Analysis of medullary B-cell density in the lipomatosis affected versus normal medullary areas of the same LNs used in the *LTB* analysis did however not show a difference (Figure 5C-D). These data support that downregulation of *LTB* can be one contributing factor to the transdifferentiation of fibroblasts at this but downregulation of *LTB* is not caused by reduced overall presence of B-cells at this site. It can also be noted that MedRCs (Nr4a1^+^ SCs and Inmt^+^ SCs) have low expression of *LTB* compared to TRCs and other BP3^+^ LN RCs (Figure EV6B), which may confer higher dependence on LTB supplied from other cells in the LN microenvironment.

### Loss of the medullary sinusoidal lymphatic network in lipomatosis with compensatory establishment of collecting-like vessels inside affected areas

In LNs with increasing degree of lipomatosis, a progressive loss of lymphatic medullary sinus area was evident with a negative correlation between the medullary sinus area and the percentage of lipomatosis (r=(-0.213)) (Figure 6A). A loss of the medullary sinus network will be expected to block the flow of lymph through the LN. Conspicuously, in LNs with more extensive lipomatosis we observed the presence of lymphatic vessels traversing the adipose tissue in the medulla, defined by expression of CLDN5 and high expression of PDPN but with low/no detection of CCL21 (Figure 6B-C). To further evaluate which molecular features define the PDPN^+^ lipomatosis associated lymphatic vessels, we stained for CD36, which we have demonstrated is a specific marker for PTX3 paracortical lymphatic vessels [11], and for MARCO which defines the medullary sinuses [11]. Lipomatosis associated vessels were negative for CD36 and displayed only occasional and low expression of MARCO (Supplementary Figure 6A-B). The vessels also rarely/or very weakly displayed LYVE-1, a marker associated with medullary sinuses (Supplementary Figure 6). The absence of CCL21, a marker of capillary initial lymphatic vessels, CD36 and low or absent MARCO and LYVE-1 together with high expression of PDPN support that the lipomatosis associated vessels likely are collecting lymphatic vessels.

**Figure 6:**
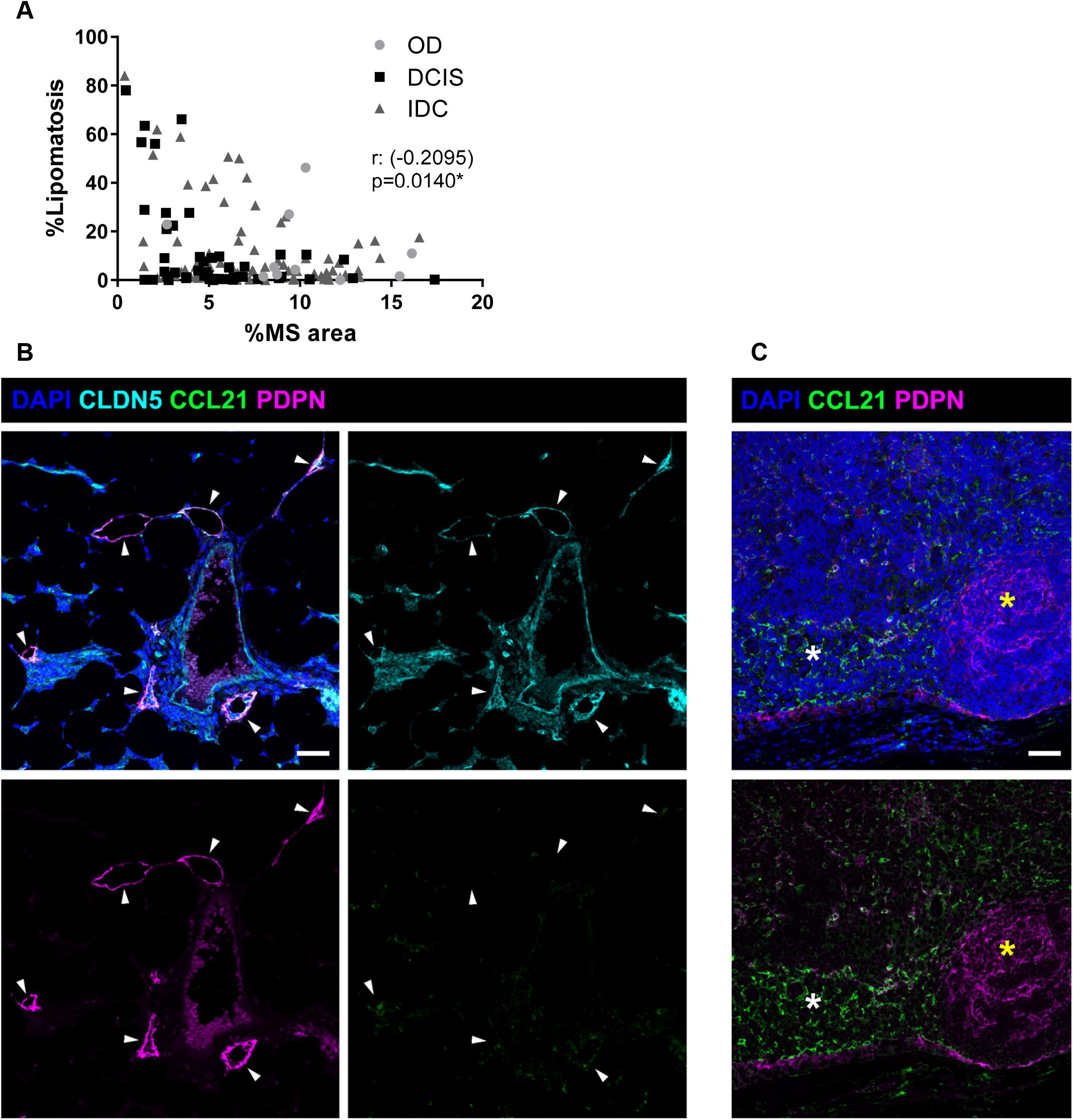
Loss of the medullary sinusoidal lymphatic network in lipomatosis with compensatory establishment of collecting-like vessels inside affected areas. **(A)** Correlation analysis of lipomatosis burden vs. percentage of medullary sinus (MS) area. n=10 lymph nodes (LNs) from n=6 organ donors (ODs), n=47 LNs from n=19 patients with ductal carcinoma in situ (DCIS), n= 80 LNs from n=35 patients with invasive ductal carcinoma (IDC). Spearman correlation test was used to test for significance. *p<0.05. **(B)** Immunofluorescence staining of collecting-like vessels in the parts of the LN affected by lipomatosis. Arrowheads visualize collecting vessels marked by high podoplanin (PDPN, magenta), presence of the vascular tight junction protein Claudin-5 (CLDN5, cyan), and the lack of the chemokine CCL21 (green). Scale bar: 50μm. **(C)** Same LN and staining as in (B) visualizing the presence of the chemokine CCL21 in the T-cell zone reticular cells (TRCs, white asterisk) and PDPN in the follicular dendritic cells (FDCs) of the B-cell zone (yellow asterisk). Scale bar: 50μm. Representative pictures are shown from ten analyzed LNs.

### Lipomatosis causes remodeling of the HEVs

Based on the dramatic loss of lymphatic vessel area and on the known potential effects of adipocytes on vascular homeostasis and inflammation [27], we next analyzed potential effects on the blood vasculature with focus on the HEVs. HEVs are characterized by their cuboidal endothelial cells and the expression of PNAd that can be detected by the MECA-79 antibody [14]. We previously showed that patients with invasive breast cancer display extensive remodeling of HEVs in the paracortex (T-cell zone) of tumor draining LNs [19] and applied the same criteria to assess HEV remodeling in lipomatosis, i.e. vessel dilation, thinning of the endothelial cells and loss of PNAd expression [19]. To avoid cancer-induced effects on HEVs in invasive breast cancer, only patients with non-invasive breast cancer, i.e. DCIS, were included in the analysis. When analyzing the difference between HEV dilation in HEVs more than 200μm from the lipomatosis, and the lipomatosis associated parenchyma (LAP), within 200μm from lipomatosis, we detected a clear difference between the locations with significantly higher proportion of dilated HEVs close to adipocytes (Figure 7A-B). Similar results were observed for thinning of the HEV endothelium and for PNAd loss with higher proportion of remodeled vessels close to adipose tissue (Figure 7A, 7C-D). A positive correlation was found between HEV dilation and thinning of the HEV endothelium as well as for HEV dilation and loss of PNAd expression, suggesting that these vascular changes are linked (Figure 7E-F). Although extensive local effects were seen on HEVs in association to parenchymal adipocytes, the extent of LN lipomatosis did not correlate with overall paracortical HEV remodeling in the patients, supporting the notion that LN lipomatosis gives local effects on HEV functions (Supplementary Figure 7).

**Figure 7:**
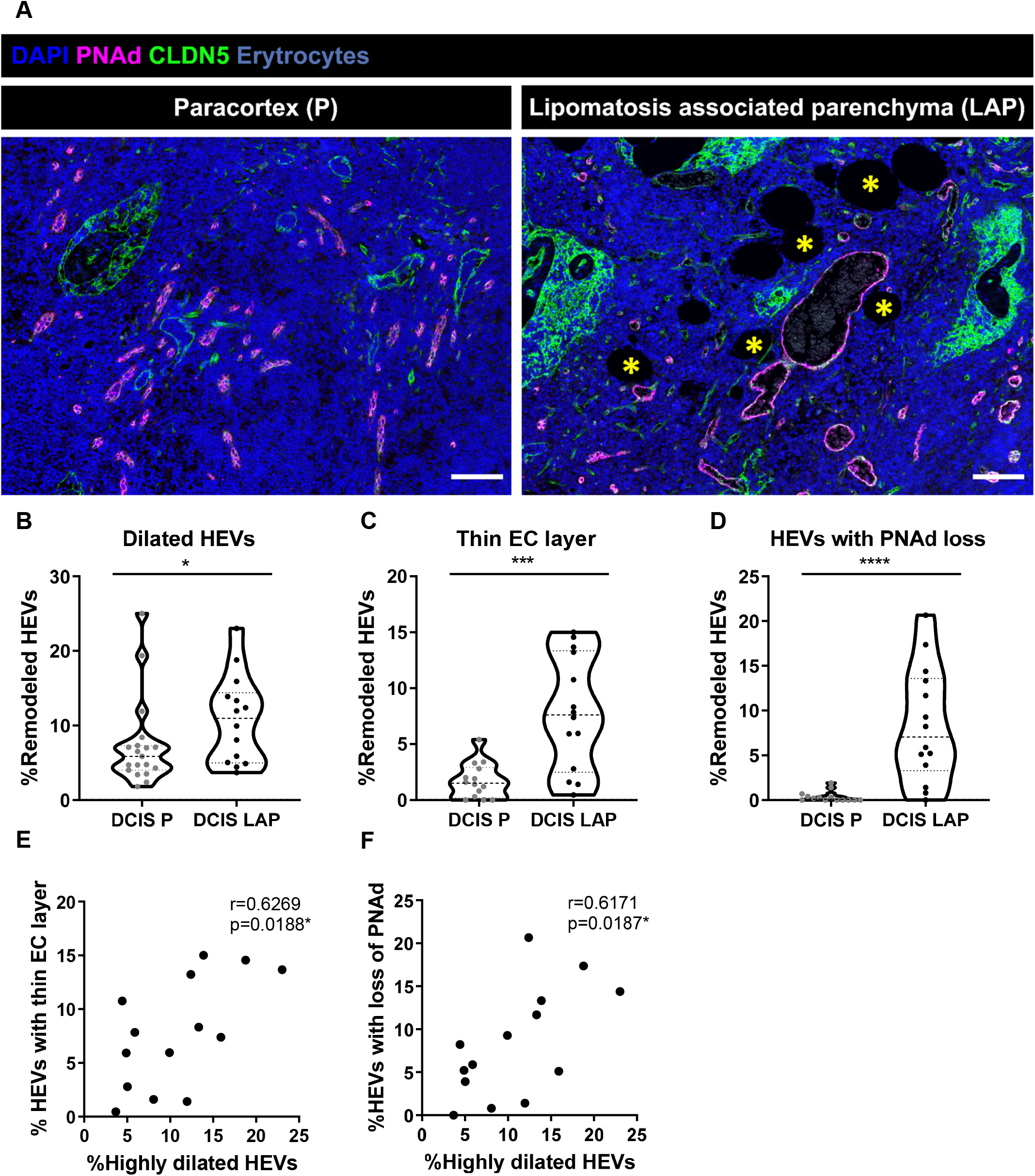
Lipomatosis causes remodeling of the high endothelial venules. **(A)** Immunofluorescence staining of paracortical (P) and lipomatosis associated parenchymal (LAP) high endothelial venules (HEVs) marked by PNAd (magenta) and Claudin-5 (CLDN5, green). Infiltrated adipocytes are marked by yellow asterisks. **(B-D)** Percentage of highly dilated HEVs, lumen>10μm **(B)**, HEVs with thin endothelial cell (EC) layer **(C)** and HEVs with loss of PNAd **(D). (E-F)** Correlation analysis of the percentage of LAP HEVs with highly dilation vs. thin EC layer **(E)** and LAP HEVs with high dilation vs. loss of PNAd **(F)**. n=19/14 lymph nodes (LNs) from 19/14 patients (P/LAP) in (A), n=14 LNs from 14 patients in (C-F). Unpaired t-test and Mann Whitney test were used for statistical analysis of P and LAP HEV remodeling, and Pearson correlation test was used for correlation analysis. *p<0.05, **p<0.01, ***p<0.001, ****p<0.0001.

### Changes in the immune contexture in LNs with lipomatosis

Naïve T-cells are continuously recruited through HEVs from blood and distribute across the paracortex guided by CCL21 expression from the surrounding TRCs . To evaluate if lipomatosis associated remodeled HEVs results in a dysfunction in T-cell recruitment we stained for the nuclear marker TCF1/7, which is constitutively expressed by naïve T-cells, memory cells and rare and long lived CD8^+^ progenitor cells [28]. Staining for TCF1/7 revealed a lower density of TCF1/7 positive cells in the immediate surrounding of highly dilated HEVs close to areas with lipomatosis compared to HEVs in the normal paracortex (Figure 8A-B). This supports that HEV remodeling induced by lipomatosis interfere with lymphocyte recruitment and/or distribution within the LN parenchyma.

**Figure 8:**
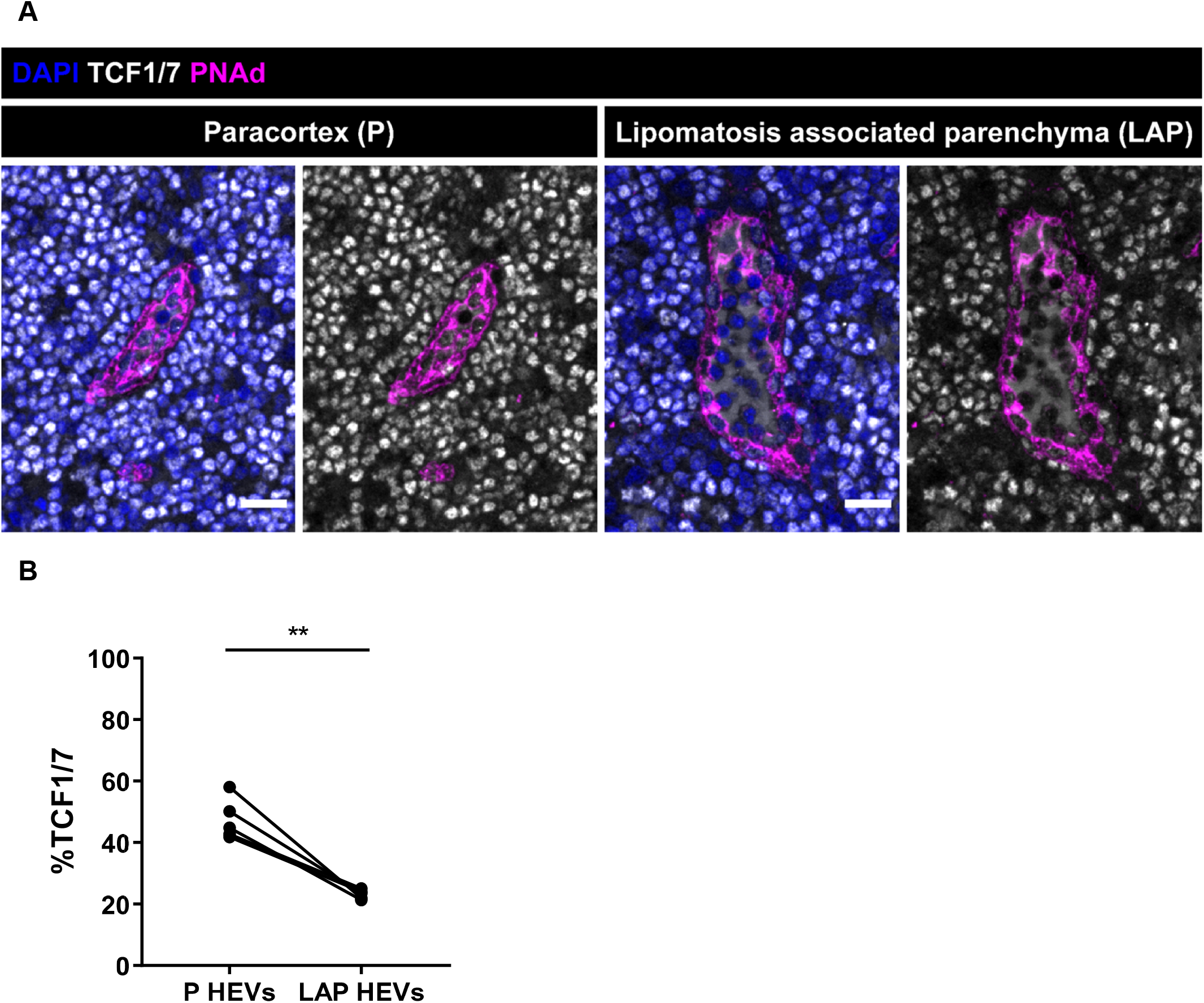
Changes in the immune contexture in LNs with lipomatosis. **(A)** Immunofluorescent staining of high endothelial venules (HEVs, PNAd) and naïve lymphocytes (TCF1/7) in lymph nodes (LNs) with intermediate to low lipomatosis, comparing areas of the paracortex (P) and the lipomatosis associated parenchyma (LAP) within the same LN. Images are representative from five LNs draining ductal carcinoma in situ (DCIS). **(B)** Paired, intra-LN analysis of the density of naïve lymphocytes in the immediate surrounding area (a layer of 2-3 immune cells) of HEVs in the paracortex and in the LAP. Dots represent the mean value of the TCF1/7 density from five different patients with DCIS. Lines represents the paired values from the same LN. Statistical analysis was performed using paired t-test. **p<0.01.

As the lipomatosis progresses the medullary cords and sinuses are replaced by adipocytes which eventually lead to loss of all normal tissue in this area (Figure 1A, 2B). The normal medulla of the LN is characterized by the presence of a dense network of macrophages and stationary plasma cells [29,30]. To evaluate possible effects on the immune cell environment in the medulla in progression of lipomatosis we staining for the macrophage marker CD68. Medullary areas with early and intermediate lipomatosis did not show any overt changes in macrophage distribution compared to unaffected areas (Supplementary Figure 8). However, analysis of plasma cells, defined by the marker CD38, showed abnormal clustering of cells within the adipose tissue (Supplementary Figure 9). In normal medulla, the plasma cells are scattered across the tissue (Supplementary Figure 9).

## Discussion

Although LN lipomatosis is a common phenomenon in the elderly, there are only a few published articles on LN lipomatosis [5,6], and both the underlying mechanisms and consequences are unknown. One reason for the lack of mechanistic data is that spontaneous LN lipomatosis rarely is seen in aging mice, where fibrosis instead dominates ([31,32]. We here choose to approach the questions behind the pathology by *in situ* analysis of human LNs.

By analyzing LNs with different degree of lipomatosis we show that it starts from deeper parts of the medullary parenchyma and demonstrate the presence of cells that display a transitional phenotype with both fibroblast and adipocyte lineage marker expression. This suggests that LN lipomatosis is driven by transdifferentiation of medullary fibroblasts into adipocytes. That LN mesenchymal cells can have an ability to transdifferentiate into adipocytes has previously been demonstrated in experimental systems *in vivo* [18,25] and for human LN-derived fibroblast *in vitro* [33]. We show for the first time that specific subsets of LN fibroblasts, i.e. MedRCs and CD34^+^ SCs, display an inherent higher sensitivity to undergo transdifferentiation into adipocytes *ex vivo*, compared to BP3^+^ RC, (i.e. TRCs, FDCs and MRCs). This provides one explanation why lipomatosis starts in the medullary regions of the LN. Although CD34^+^ SCs displayed the strongest sensitivity, these cells mainly build up the capsule of the LNs and are only found around adventitia of large blood vessels inside the LN. At later stages of lipomatosis it is likely that other subsets including TRCs may contribute, possible driven by advancing disruption of homeostasis.

It has previously been demonstrated that LTBR signaling can counteract the differentiation of LN stromal cells into adipocytes in mouse experimental systems *in vivo* [18,25]. A common progenitor for LN fibroblasts and adipocytes was also identified in early development of the LN anlage [18]. In this context, it is interesting to note that we could confirm reduced expression of *LTB* in areas affected by lipomatosis, suggesting this could be one contributing factor driving transdifferentiation of fibroblasts into adipocytes in the aging LN. Our *in silico* analysis of mouse MedRCs also support that these cells express much less *LTB* than TRCs, which may make them more sensitive to loss of LTB from other cells in the microenvironment. While B-cells are known to be a major source of LTB [26], we did not detect a reduction in B-cell density. Thus, functional changes of medullary B-cells and/or other cell types may contribute to downregulation of *LTB*. Based on our data it is likely that combinatorial effects driven by microenvironmental changes such as downregulation of *LTB*, together with a higher sensitivity for MedRC and potentially also rare CD34^+^ SCs in these areas, drives initiation and progression of lipomatosis. In addition to LTBR signaling the Hippo pathway has been implicated in regulation of LN stromal cell differentiation [25]. *In silico* analysis of the dataset published by Rodda *et al*., indeed confirmed enrichment of genes associated both with NF-kappa B signaling, and the Hippo pathway in TRCs, FDCs and MRCs when compared to the MedRCs and CD34^+^ SCs.

As part of the phenotype in lipomatosis affected LNs, we observed a dramatic reduction of lymphatic medullary sinus area and the medullary lymphatic endothelium is progressively lost. Our data do not support that endothelial cells transform into adipocytes like the fibroblasts, but loss of ECM interactions which are dependent on surrounding fibroblast, may lead to apoptosis of LECs and contribute to the observed pathology. Experimental system able to mimic the pathology in the mouse would however be required to evaluate this hypothesis. A degeneration of the LN lymphatic sinuses is expected to result in obstruction of lymph flow. We however observed the appearance of collecting-like lymphatic vessels inside the areas of the LN affected by lipomatosis, which may explain why LN lipomatosis can progress and affect multiple LNs without causing lymph edema in the patients. These vessels were defined by higher expression of PDPN than most other LN lymphatic vessels, lacked CD36 (marker expressed by paracortical sinuses in the LN [11]), displayed low or absent MARCO and LYVE-1, both markers of medullary sinuses. They also have no detectable expression of CCL21, which is a marker highly expressed by initial capillary lymphatic vessels in the periphery. It is interesting to speculate that these vessels result from ingrowth of collecting vessels through the destructed areas of the LN.

Besides the striking effect on the lymphatic vascular medullary sinus area, we show that lipomatosis in the analyzed human LNs also cause extensive local remodeling of HEVs. Loss of LTBR signaling has been linked to dysregulation of HEVs in mouse models [34,35] and the downregulation of *LTB* in affected areas we observe is a likely contributing factor, but we cannot exclude additional cooperating mechanisms. Adipocytes express high levels of a number of growth factors [36], including vascular endothelial growth factor A (VEGFA), which potentially also could contribute to disturbing HEV homeostasis. That the effects caused by lipomatosis are local is emphasized by the lack of correlation to paracortical HEV remodeling, a phenomenon we previously have shown is common in tumor draining LNs [19]. Notably, we here also show that the remodeling of HEVs in lipomatosis is linked to reduced density of TCF1/7 naïve T-cells around dilated HEVs. Thus, already early and intermediate lipomatosis have implications for the immune contexture of the human LN. Consistent with this notion, we observed accumulation of CD38 plasma cells within the adipose tissue. CD38+ plasma cells are normally scattered across the medullary cords and the clusters of cells seen in areas with adipocytes indicates that the cells are trapped in the adipose tissue. As lipomatosis progresses, the normal LN tissue, including the immune cells, medullary cords and sinuses, is replaced by adipocytes, which is likely to in late stages, cause a complete loss of its important immunogenic functions.

Already in early stages, LN lipomatosis is likely to affect both lymphocyte recruitment into the LN through HEVs and recirculation of lymphocytes back to the blood through the lymphatic sinuses; which are both fundamental functions for induction of adaptive immunity [8]. Consequently, LN lipomatosis is highly likely to have impact on vaccination in the elderly by reducing access to functional lymphoid tissues. Clinical studies in elderly patients with different degrees of LN lipomatosis are however needed to confirm this hypothesis. Of interest in this context, there are indications from older literature that LNs in different draining areas are affected to different degrees [5], why it can be speculated that the site for administration of vaccination may be a way to circumvent the effects caused.

From a perspective of cancer development adipose tissue has gained a lot of attention [37]. Adipocytes can contribute to a disturbed microenvironment promoting tumor formation and have been speculated to contribute to a pre-metastatic niche in for example the bone marrow [38]. The bone marrow, similar to the LN, display infiltration of adipocytes with age and consequently display reduced output of hematopoietic stem cells [38,39]. It cannot be excluded that LN lipomatosis contribute to a pre-metastatic niche in the context of nodal metastasis, thus patients with high degree of LN lipomatosis may be more likely to develop growth of metastasis inside the LN. To evaluate this in human cancer would be of high interest for future studies, but is out of the scope for this paper. Since the phenomenon is so common in general aging, it would require careful examination of LN lipomatosis and metastasis in all axillary LNs across very large groups of patients.

The loss of LN parenchyma due to lipomatosis is highly likely to interfere with induction of adaptive immunity and is likely to contribute to a reduced immune status of the aging induvidual. This may have multitude influences on the ability to respond to infections and vaccination as well as diminished ability to mount anti-tumor responses in cancer highlighting a need for further research.

## Materials and methods

### Biobank material and ethical considerations

Formalin fixed and paraffin embedded (FFPE) biobank LNs from breast cancer patients were used for immunostaining and RNAscope. The patient cohort includes patients with the noninvasive DCIS (patients: n=22, LNs: n=65) and the invasive IDC without LN metastasis (patients: n=44, LNs: n=115). Pancreatic LNs from organ donors (ODs) were FFPE and used as controls (donors: n=14, LNs: n=66). The approval to use biobank and OD LNs was provided by Uppsala regional ethics committee, 2017/061 and addition 2017/061:1 and 2017/061:2 to Ulvmar MH. Mice for tissue isolation were held under ethical permit 6009/17 approved by the Uppsala Animal Experiment Ethics Board to Ulvmar M.H.

### Immunofluorescence staining

The protocol for the immunostaining was previously described (Bekkhus *et al*. [17]). In short; FFPE LNs were treated with xylene and ethanol gradient to remove paraffin and rehydrate the tissues. Antigens were retrieved in 1mM ethylenediaminetetraacetic acid (EDTA) (pH9) (Invitrogen) at 97°C for 20min. The tissues were blocked with 0.033% Streptavidin (Sigma), 0.0033% Biotin (Sigma), in the case of biotin-streptavidin amplification, and 5% donkey serum (Sigma), all diluted in PBS with 0.05% Tween20 (Merck) (PBST), for 15, 15 and 20min respectively. The tissues incubated in primary antibodies (Supplementary Table 1) at 4°C overnight. The tissues were washed in PBST and incubated in secondary antibodies (Supplementary Table 2) for 30min at room temperature. For streptavidin-biotin amplification, the tissues also incubated in CF647 conjugated streptavidin (Biotium) for 30min at room temperature. For nuclear counterstaining the tissues incubated in 4′,6-diamidino-2-phenylindole (DAPI) (Invitrogen) for 5min and the slides were mounted with ProLongGold (Invitrogen) and #1.5 coverslips.

### Cell sorting for adipogenic culture

Inguinal, brachial and axillary LNs (n=210-240 per experiment) were harvested and digested as described by Xiang *et al*. [11]. CD45^+^ immune cells were depleted from the samples using CD45 beads and LS-columns following the manufacturer’s instructions (Miltenyi Biotec). The pellets were resuspended in Fc block (1:100 in FACS buffer containing EDTA and fetal bovine serum (FBS) 0.5%) and incubated shortly before adding primary antibodies that incubated on ice for 30min (Supplementary Table 3). The cells were washed by adding FACS buffer. Sytox Blue (Invitrogen) dead cell staining was added before the sorting. Cells were sorted on a BD FACSAria III (BD Biosciences) (100 um nozzle, 20 pounds per square inch (psi), and an acquisition rate of 500–2,000 events per second). Shortly, live single cells were gated using FSC-A/SSC-A followed by FSC-H/FSC-W and SSC-H/SSC-W. LN fibroblasts (CD45^neg^, CD11b^neg^, PECAM1^neg^, PDPN^pos.^) were sorted into three fractions based on the markers CD34 and BP3 (Bst1): BP3^neg^ CD34^neg^ MedRCs, BP3^neg^ CD34^pos^ CD34^+^ SCs and BP3^pos.^ CD34^neg^ RCs BP3^neg^ CD34^neg^ MedRCs, BP3^neg^ CD34^pos^ CD34^+^ SCs and BP3^pos.^ CD34^neg^ RCs (i.e. TRCs, FDCs and MRCs). 60000-110000 cells were collected per population and transferred for culture in coverslip covered petri dishes with StableCell™ MEM, alpha modification media (Sigma Aldrich) supplemented with 1% L-glutamine (Invitrogen), 1% Penicillin-Streptomycin, 10% FBS and anti-LTBR antibody (1:200, ab65089, Abcam). After 3 days of culture, control cells were fixed and treatment with adipogenic media (0.5mM IBMX, 10μg/ml Insulin, and 1μM dexamethasone and 100μM indomethasine) was induced and continued for 3 days before fixing in 4% PFA and staining for Perilipin and αSMA-AF647 (Supplementary Table 1 and 3). For nuclear counterstaining, the cells incubated in DAPI (Invitrogen) for 5min.

### RNAscope

The RNAscope experiment was performed according to the manufacturer’s instructions (RNAscope® Fluorescent Multiplex Kit User Manual Part 1 & 2, Advanced Cell Diagnostics, INC.) using their kit for amplification. In short, FFPE LNs were sectioned at 4μm under RNase free conditions. Sections were melted for 1h at 60°C and treated with xylene and absolute ethanol to remove paraffin. Hydrogen peroxidase was blocked by incubation with hydrogen peroxide for 10min and antigens were retrieved by incubation in 1X RNAscope Target Retrieval Reagent for 15min at 99°C. Sections were treated with RNAscope Protease Plus and incubated with probes for *LTB* (310471-C2) which were left to hybridize for 2h at 40°C. Sections were stored in 5X Saline Sodium Citrate (SSC) buffer overnight at RT. Next, Amp1, Amp2 and Amp3 were left to hybridize for 30, 30 and 15min respectively at 40°C. The HRP-C2 signal was developed by incubation with HRP-C2 for 15min at 40°C followed by Opal570 (1:1500) for 30min at 40°C and HRP block for 15min at 40°C. The slides were counterstained with DAPI and mounted with ProLongGold and #1.5 coverslips. Tissues were all imaged the next day.

### Imaging

For image acquiring of immunofluorescent stainings of sections, the Vectra Polaris™ Automated Quantitative Pathology Imaging System (Akoya Biosciences) was used. For the imaging of RNAscope and cell cultures, the LSM 700 confocal microscope (Zeiss) with Plan-Apochromat 63x/1.40 Oil DIC M27 and Plan-Apochromat 20x/0.8 M27 objectives were used respectively.

### Image analysis

All image analysis was performed using QuPath 0.2.3, Fiji Image J version 1.52p. and Phenochart 1.0.12 (Akoya Biosciences).

The presence of lipomatosis was scored based on the presence of adipocytes within the capsule in all LNs from all patients/donors to give each patient/donor a status of being affected by lipomatosis or not.

The lipomatosis burden was quantified as a percentage of the total LN area using QuPath and Fiji Image J. The channel for DAPI and the autofluorescence channels AF488 and/or AF555 were used to make a region of interest (ROIs), defining the intracapsular LN parenchyma as the total LN area, by using the “wand” tool in QuPath. The ROIs were exported to Fiji Image J. To define the lipomatosis area, a threshold was set on the DAPI channel to cover the adipocytes with limited background and the threshold was converted to a binary mask with dark background. A threshold without dark background was set on the binary mask and the ROI for the LN area was used to measure the lipomatosis area. To limit the detection of non-adipocytic area, separate ROIs were made for the adipocytes to measure the lipomatosis area in LNs with a lot of background (less dense network of DAPI). LNs with no morphologically confirmed adipocytes were set to 0% lipomatosis and LNs with less than three adipocytes were excluded.

The localization of the lipomatosis was defined based on the pattern of the DAPI, CLDN5, CCL21 and PNAd staining. The cortex was defined by the dense DAPI staining (B-cell follicles), absence of CCL21^high^ fibroblasts and PNAd and/or CLDN5 positive vessels. The paracortex was defined by CCL21^high^ fibroblasts and PNAd/CLDN5 positive vessels. The medulla was defined by the presence of CLDN5 positive lymphatic sinuses and the absence of CCL21^high^ fibroblasts. All LNs were in the end categorized individually based on the localization of the lipomatosis; medulla, medulla/paracortex, or medulla/paracortex/cortex. The analysis was performed using Phenochart. LNs with less than three adipocytes were excluded from the analysis.

To analyze if Perilipin was expressed by fibroblasts or LECs, the localization was first determined based on the staining of Perilipin, LYVE-1, αSMA and CD19. LYVE-1 marked the medullary sinuses, αSMA the fibroblasts of the medulla and paracortex and CD19 marked the B-cell follicles and the B-cells of the medulla. If the Perilipin^+^ adipocytes were present in fibroblastic compartment they were checked for co-staining with αSMA. If they were present within the medullary lymphatic sinuses, they were first checked for co-staining with LYVE-1 but due to LYVE-1 having a high heterogeneity in the human medullary lymphatic sinuses these LNs were also stained for CLDN5 to rule out co-expression with the medullary lymphatic sinus LECs. For the vector based localization analysis in Supplementary Figure 1, the “Plot Profile” tool in Fiji Image J was applied on a vector drawn across the outline of the αSMA^+^/Perilipin^+^ cell.

The presence of transformed fibroblasts from the FACS sorted cultures was quantified by using Fiji image J. A threshold was set on the DAPI channel (7-255) to mark all the nuclei. The “Analyze Particles” tool was applied and the nuclei were added to the ROI manager, excluding particles on the edges. If several nuclei were assigned to one ROI, separate ROIs were manually added to get the correct number of ROIs corresponding to the staining. The ROIs were then applied onto the Perilipin channel where a threshold was set (10-255). Each nuclear ROI was assigned as being Perilipin positive or negative in the immediate surrounding area. Eight 20x frames per subset were analyzed containing a mean number of 118 cells per frame and the percentage of Perilipin positive cells was calculated.

RNAscope images were analyzed using Fiji Image J. Samples with low to intermediate lipomatosis were selected to allow analysis of medullary regions with and without adipocytes within the same LN. Five 63x frames were analyzed per LN and per condition (medulla far from adipocytes vs. medulla close to adipocytes). Thresholds for DAPI and LTB were set and the areas were measured. Due to high patient and RNA quality variations as a consequence of using human biobank samples, a paired analysis was performed within the same LN, comparing the medullary areas far from and close to adipocytes, and the thresholds for DAPI and LTB were adjusted for the specific sample.

The B-cell density analysis was performed by using Fiji Image J. The same LNs as in the RNAscope analysis were stained for CD19 and CLDN5. Six regions of interests were selected based on the areas used for analysis of the RNAscope sections, three far from adipocytes and three close to adipocytes. Areas occupied by medullary lymphatic sinuses (CLDN5^+^ staining) and adipocytes (morphologically defined as adipocytes with lack of staining of CLDN5, CD19 and DAPI) were excluded from the area of analysis. A threshold was set for CD19 and the percentage of CD19 area was quantified. Individual thresholds were set for different LNs due to intra LN analysis.

The percentage of medullary sinus area was quantified using Fiji Image J. The LN area was reused from the analysis of the lipomatosis burden and a threshold was set on the CLDN5 channel. A ROI was made for the region of the LN occupied by medullary and trabecular lymphatic sinuses, which transverse into the medulla of the human LN, and the percentage of medullary sinus (MS) area out of the total LN area was calculated.

The presence of collecting-like vessels in the lipomatosis affected regions was analyzed by using Phenochart. The collecting-like vessels were defined as PDPN^high^/CLDN5^+^/CCL21^-^/CD36^-^/MARCO^low-negative^/LYVE-1^low-negative^ and by being embedded in adipose tissue. Only LNs with high lipomatosis were included for analysis.

For the analysis of HEV remodeling, red green blue (RGB) snapshots of lipomatosis associated HEVs were created using Phenochart and were analyzed in Fiji Image J. The number of PNAd^+^/CLDN5^+^ vessels was quantified and vessels within 200μm from adipocytes were defined as lipomatosis associated HEVs. The lumen diameter was measured as a mean of three independent measurements of the shortest diameter to reduce effects of the angle of sectioning. Each vessel was classified as non-dilated (no lumen), intermediately dilated (0-10μm), highly dilated (>10μm) and/or having a thin endothelial cell layer or loss of PNAd expression in parts of the vessel.

TCF1/7^+^ cell presence was analyzed using Fiji Image J. Highly dilated HEVs, surrounded by CCL21^+^ fibroblasts, within 200μm from adipocytes were selected by creating a ROI for this region. Only HEVs with a clear immune cell perimeter were included to limit the bias of having an adipocyte in the frame. As controls, non-dilated HEVs, surrounded by CCL21^+^ fibroblasts, more than 200μm from adipocytes were selected by creating a ROI for this region. All HEVs were selected unbiased of the TCF1/7 staining, using a composite image of DAPI, PNAd and CCL21 only for the selection of vessels. A threshold was set for the PNAd channel and was applied to create a binary image. Background was removed by deleting pixels outside the HEV, using the freehand selections tool, and only the binary mask of the HEV was kept. If there was a loss of PNAd, the outline of the HEV was filled out by using the “paintbrush tool” focusing on the composite image where the outline of the vessel is structurally visible. The binary tool to “fill holes” was used and the mask was added as a ROI for the HEV. The binary image was dilated by 20 iterations to increase the size of the mask of the HEV using the binary tool “dilate”. Twenty iterations equals a layer of approximately 2-3 lymphocytes. The channel for TCF1/7 was selected and a threshold was set. The ROI for the HEV only and the 20x dilated ROI was used to measure the area of TCF1/7 outside of the LN by subtracting the area of the HEV itself from the 20x ROI area. The percentage of this area covered by TCF1/7 was used for data analysis. As with the RNAscope and CD19 analysis, due to high patient variations as a consequence of using human biobank samples, a paired analysis was performed within the same LN, comparing the areas far from and close to adipocytes, and the thresholds for PNAd and TCF1/7 were adjusted for the specific sample. Three highly dilated HEVs and three non-dilated HEVs were analyzed per patient including five patients from the DCIS group.

### Statistical analysis

Statistical analysis was performed with GraphPad Prism version 7 and 8. For correlation analysis, Spearman or Pearson correlation tests were used based on the distribution of the data analyzed. For comparison between normally distributed paired groups, paired t-test was used. In cases of non-paired groups, unpaired t-test or Mann Whitney test were used based on the distribution of the data analyzed. In the case of more than two groups of normally distributed data, one-way ANOVA and Holm-Sidak’s multiple comparisons test were used.

### Bioinformatical analysis

The processed single cell RNAseq data of lymph node stromal cells and cell annotation were obtained from the previous public study [10]. Cell types were stratified using Bst1gene expression. “TRC” (“Ccl19^hi^ TRC”), “Ccl19^lo^ TRC”,”Cxcl9^+^ TRC”, “MRC”,”FDC” were defined as Bst1(BP3) high population; cell types of “Nr4a1^+^ SC”, “Inmt^+^ SC” and “CD34^+^ SC” were defined as Bst1 low population. We pooled clusters with high Bst1 expression (“TRC”,”Ccl19^lo^ TRC”,”Cxcl9^+^ TRC”, “MRC”,”FDC”), and compared with clusters of “CD34^+^ SC”, “Nr4a1^+^ SC”, “Inmt^+^ SC” (expressing low Bst1), respectively. Wilcoxon Rank Sum test was applied for identifying differentially expressed genes (DEG) in Seurat package (v. 4.1.1 PMID: 25867923). For cluster “CD34^+^ SC” comparing with clusters expressing high Bst1 expression, 482 genes were identified as statistically significant expressed (Bonferroni corrected p value<0.01, fold change (log2)>0.5 or <-0.5). For cluster “Nr4a1^+^ SC” comparing with clusters expressing high Bst1 expression, 215 genes were identified as statistically significant expressed (p value and fold change were used same as describe above). For cluster “Inmt^+^ SC” comparing with clusters expressing high Bst1 expression, 274 genes were identified as statistically significant expressed (p value and fold change were used same as describe above). Heatmap visualization was produced using ComplexHeatmap package (v. 2.10.0, PMID: 27207943).

## Supporting information

all supplmental files

## Acknowledgments

The authors thank Professor Olle Korsgren, Uppsala University, for access to pancreatic lymph nodes from organ donors and Daniel Vasiliu Bacovia for selecting patients for analysis. We also thank Liqun He, Uppsala University, for initial bioinformatical analysis. This research was funded by the Swedish Research Council (2016-02492), Swedish Cancer Foundation (2017/759 and 20 0970 PjF) and Kjell and Märta Beijer Foundation to Ulvmar MH.

## Author Contributions

Conceptualization, M.H.U. and T.B; analysis and Figures, T.B., Y.S. and A.O.; bioinformatic analysis, Y.S., writing original manuscript, M.H.U. and T.B.; Editing of manuscript, all authors. Project administration and funding, M.H.U. All authors have read and agreed to the manuscript.

## The paper explained

### Problem

Lymph nodes are small organs that functions as the headquarters of our adaptive immune defense. Within the lymph nodes lymphocytes (i.e. T-cells and B-cells) become activated and get their instructions to respond to vaccination and to protect us against infections or against cancer. However, with aging the lymph nodes change, and the normal tissue is changed into fat, a pathological process called lipomatosis. Although LN lipomatosis is a very common in lymph nodes of elderly the mechanisms driving these changes and the potential effects on the lymph node immune functions have not been studied which constitute a major gap our current knowledge.

### Results

This paper show the first evidence that lipomatosis in the aging human lymph node is caused by transdifferentiation of fibroblasts into adipocytes and starts from the central part of the lymph node called the medulla. We also show that fibroblast isolated from this region display inherent higher sensitivity to transdifferentiation compared to fibroblast from the T-cell zone. Molecularly we demonstrate that lipomatosis if associated with local downregulation of a cytokine called LTB, a factor known to counteract adipogenesis. We also show that the consequences of lipomatosis include an extensive vascular remodeling with a progressive loss of the medullary lymphatic vasculature and dramatic remodeling of the specialized blood vessels within the lymph node, called high endothelial venules (HEVs). The latter can be linked to reduced density of naive TCF1/7^+^ lymphocytes around the HEVs and accumulation of CD38 plasma cells in the affected areas. Thus causing major changes to the immune microenvironment.

### Impact

Our data put focus on lymph node lipomatosis as major factor that can contribute to decreased immune functions in the elderly and both warrant an increase awareness of lymph node lipomatosis as a contributing factor in human disease and immune-directed therapy and vaccination, highlighting a need for continued research.

## Supplementary Tables and their legends

**Supplementary Table 1:** Primary antibodies. Table of primary antibodies used in the immunofluorescence staining. Including information of the antigen, host, class, clone, company and dilution. (na: non-applicable)

| Antigen: | Host: | Class: | Clone: | Company: | Dilution: |
| --- | --- | --- | --- | --- | --- |
| CCL21 | Goat | Polyclonal | na | R&D Systems, | 1:20 |
| CD19 | Rat | Monoclonal | 6OMP31 | Invotrogen | 1:100 |
| CD38 | Rabbit | Polyclonal | na | Atlas antibodies | 1:50 |
| CD68 | Mouse | Monoclonal | Kp-1 | Sigma-Aldrich | 1:50 |
| Claudin-5 | Rabbit | Monoclonal | EPR7583 | Abcam | 1:100 |
| Claudin-5 | Mouse | Monoclonal | 4C3C2 | Invtrogen | 1:50 |
| LYVE-1 | Goat | Polyclonal | na | R&D systems | 1:100 |
| PDPN | Mouse | Monoclonal | D2-40 | Dako | 1:50 |
| Perilipin | Rabbit | Polyclonal | na | Invitrogen | 1:50 |
| PNAd | Rat | Monoclonal | MECA-79 | BD Pharmingen | 1:20 |
| TCF1/7 | Rabbit | Monoclonal | C63D9 | Cell Signaling | 1:50 |
| αSMA | Mouse | Monoclonal | 1A4 | Dako | 1:100 |
| αSMA-AF647 | Mouse | Monoclonal | 1A4 | Santa Cruz Biotechnology | 1:20 |

**Supplementary Table 2:** Secondary antibodies. Table of secondary antibodies used in the immunofluorescence staining. Including information of species reactivity, conjugate, catalogue number, company, and dilution.

| **Species Reactivity:** | **Conjugate:** | **Company:** | **Dilution:** |
| --- | --- | --- | --- |
| Goat | AF488 | Invitrogen | 1:300 |
| Goat | AF555 | Invitrogen | 1:300 |
| Goat | AF594 | Invitrogen | 1:300 |
| Mouse | AF488 | Jackson ImmunoResearch | 1:300 |
| Mouse | Cy3 | Jackson ImmunoResearch | 1:300 |
| Mouse | AF647 | Jackson ImmunoResearch | 1:300 |
| Rabbit | AF555 | Invitrogen | 1:300 |
| Rabbit | AF594 | Invitrogen | 1:300 |
| Rabbit | AF647 | Invitrogen | 1:300 |
| Rat | AF488 | Jackson ImmunoResearch | 1:300 |
| Rat | AF647 | Jackson ImmunoResearch | 1:300 |
| Rat | Biotin | Jackson ImmunoResearch | 1:200 |

**Supplementary Table 3:** Primary antibodies for FACS. Table of primary antibodies used in the FACS staining. Including information of the antigen, host, class, clone, company, dilution and conjugate.

| **Antigen:** | **Host:** | **Class:** | **Clone:** | **Company:** | **Dilution:** | **Conjugate:** |
| --- | --- | --- | --- | --- | --- | --- |
| BP3 | Mouse | Monoclonal | BP-3 | Biolegend | 01:50 | APC |
| CD11b | Rat | Monoclonal | M1/70 | Invitrogen | 01:50 | PerCP-Cy5.5 |
| CD31 | Rat | Monoclonal | 390 | eBioscience | 01:50 | PE-Cy7 |
| CD34 | Rat | Monoclonal | RAM34 | Invitrogen | 01:50 | FITC |
| CD45 | Rat | Monoclonal | 30-F11 | Invitrogen | 01:50 | PerCP-Cy5.5 |
| PDPN | Syrian Hamster | Monoclonal | eBio8.1.1 | eBioscience | 01:50 | PE |
| Ter119 | Rat | Monoclonal | TER-119 | Biolegend | 01:50 | BV421 |

## Figure legends for supplementary Figures

**Supplementary Figure 1: Image analysis of αSMA and Perilipin signal localization in a lipomatosis affected lymph node. (A-B)** Immunofluorescence images of αSMA **(A)** and Perilipin **(B)** with vector shown in white; size: 4.6μm. **(C-D)** Vector based profile plot of αsma **(C)** and Perilipin **(D)**.

**Supplementary Figure 2: Visualization of different fibroblast subsets of the human lymph node**. Immunofluorescence staining of B-cells (CD19; blue) marking the cortex (C) and part of medulla (M), the chemokine CCL21 (magenta), marking the fibroblastic reticular cells also known as T-cell reticular cells (TRCs) of the paracortex (P), and alpha smooth muscle actin (αSMA; green) marking both the TRCs of the paracortex (P) and the medullary reticular cells (MedRC) in the medulla (M). MedRCs are low or negative for CCL21, in contrast to TRCs that express high levels. Scale bar: 150μm.

**Supplementary Figure 3: Expression pattern of CD34 in the human lymph node**. Immuno-fluorescent staining of a lymph node with low lipomatosis stained for CD34 (cyan), CCL21 (staining the paracortex, magenta) and DAPI (blue). CD34 is expressed in the fibroblasts of the capsule (Cap) and its invaginations called trabecula found in the trabecular sinuses (TS) (red asterisks) and in the adventitia of larger blood vessels (yellow asterisks). It is also expressed by capillary blood vessels of the germinal center (GC) of the cortex (C) and medulla (M) (green asterisks), and by the high endothelial venules (HEVs) of the paracortex (P) (white asterisks). Scale bar 250μm.

**Supplementary Figure 4: FACS**. LN fibroblasts (CD45^neg^, CD11b^neg^, PECAM1^neg^, PDPN^pos.^) were sorted into three fractions based on the markers CD34 and BP3 (Bst1): BP3^neg^ CD34^neg^ MedRCs, BP3^neg^ CD34^pos^ CD34^+^ SCs and BP3^pos.^ CD34^neg^ RCs BP3^neg^ CD34^neg^ MedRCs, BP3^neg^ CD34^pos^ CD34^+^ SCs and BP3^pos.^ CD34^neg^ RCs (i.e. TRCs, FDCs and MRCs). Dot plots display gating strategy and sorted populations are indicated.

**Supplementary Figure 5: Expression analysis. (A)** Bar plot displaying expression of *Ltbr* for each cell in the different clusters described by Rodda et al., (10). (**B**) Heatmap of DEG genes (“CD34^+^ SC” cluster expressing low Bst1 vs. cell clusters expressing high Bst1) in the NFkap-paB pathway, differently expressed in subsets described by Rodda et al., Values are normalized counts (log-scaled). (**C**) Heamap of DEG genes (“CD34^+^ SC” cluster expressing low Bst1 vs. cell clusters expressing high Bst1) in the Hippo pathway, differently expressed in subsets described by Rodda *et al*., Values are normalized counts (log-scaled). The MedRCs terminology, used in other parts of this paper includes the two subsets Nr4a1+ SC and Inmt+ SC, which are both BP3^neg/low^ CD34^neg^, the three TRC subsets, FDCs and MRCs are all BP3 high subsets, while CD34+ SC are BP3 low/neg.

**Supplementary Figure 6. CD36, MARCO and LYVE-1 expression in collecting-like vessels**. Immunofluorescence images of lymph nodes (LNs) with intermediate to high lipomatosis. **(A)** LN stained for CD36 (green) and podoplanin (PDPN, magenta). **(B)** LN stained for MARCO (green) and PDPN (magenta). **(C)** LN stained for LYVE-1 (green) and PDPN (magenta). White arrowheads points out collecting-like vessels. Scale bar: 50μm.

**Supplementary Figure 7: Lipomatosis is inducing a local remodeling of the high endothelial venules**. Correlation analysis of the percentage of highly dilated high endothelial venules (HEVs) in the paracortex (more than 200μm from the adipocytes) vs. the percentage of lipomatosis in the lymph node (n=19 patients). Spearman correlation test was used for statistical analysis. No correlation was detected; r=0.01888, p=0.9388.

**Supplementary Figure 8: Macrophage distribution in a lipomatosis affected lymph nodes**. Immunofluorescent staining of lymph nodes with low and intermediate lipomatosis stained for the macrophage marker CD68 (green), Claudin-5 (CDLN5, magenta), CD19 (white) and DAPI (blue). Scale bar: 100μm.

**Supplementary Figure 9: Distribution of CD38 plasma cells in the medulla of lipomatosis affected lymph nodes**. Immunofluorescent staining of CD38 (cyan), Claudin-5 (CLDN5, green), CD19 (magenta) and DAPI in areas of a lymph node (LN) with different degree of lipomatosis. Scale bar: 200μm. Pictures representative from nine LNs.

