## Supplementary material for "Stromal transdifferentiation drives lymph node lipomatosis and induces extensive vascular remodeling": all supplmental files

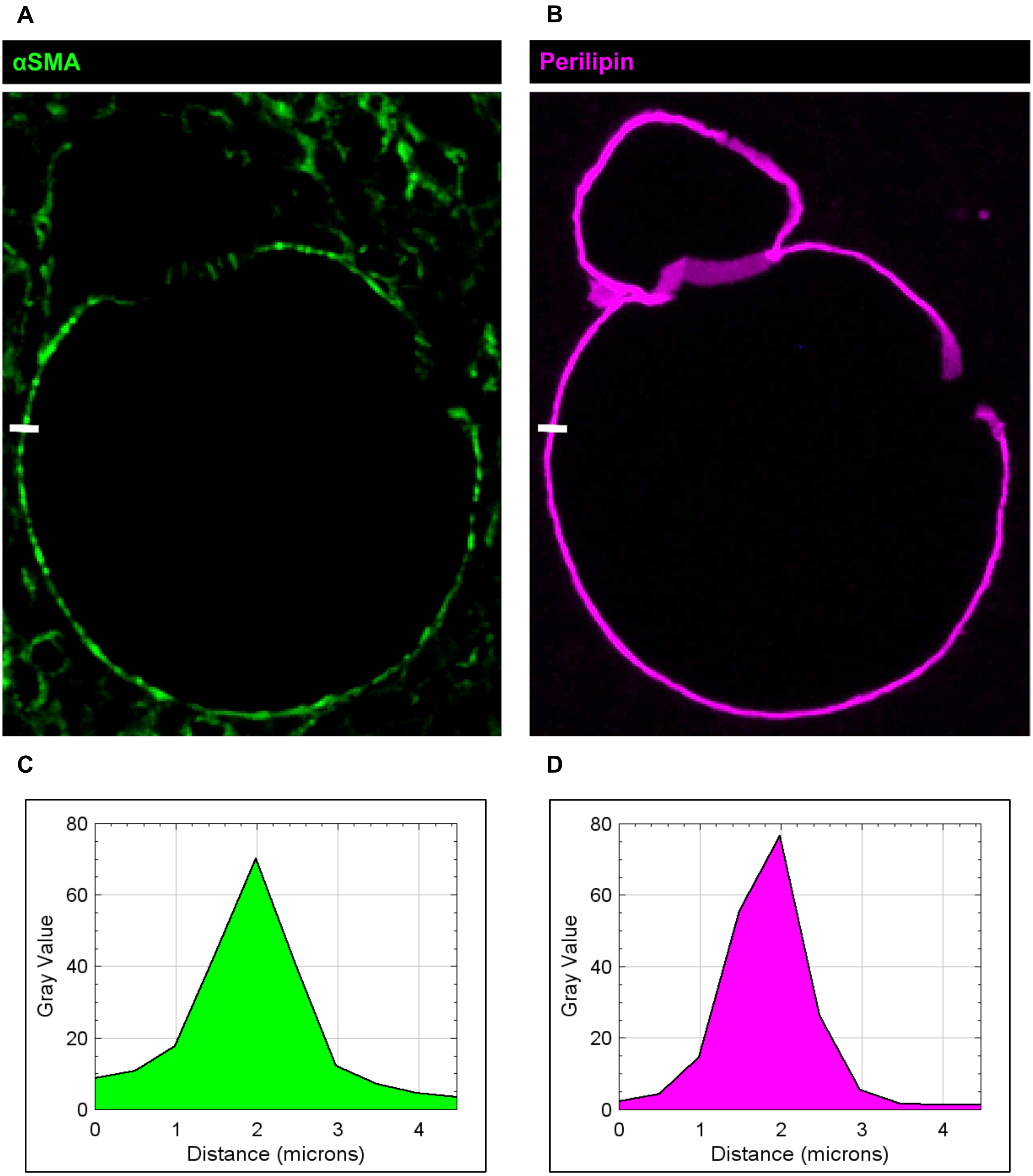


**Supplementary Figure 1: Image analysis of αSMA and Perilipin signal localization in a lipomatosis affected lymph node. (A-B)** Immunofluorescence images of αSMA **(A)** and Perilipin **(B)** with vector shown in white; size: 4.6µm. **(C-D)** Vector based profile plot of αSMA **(C)** and Perilipin **(D).**

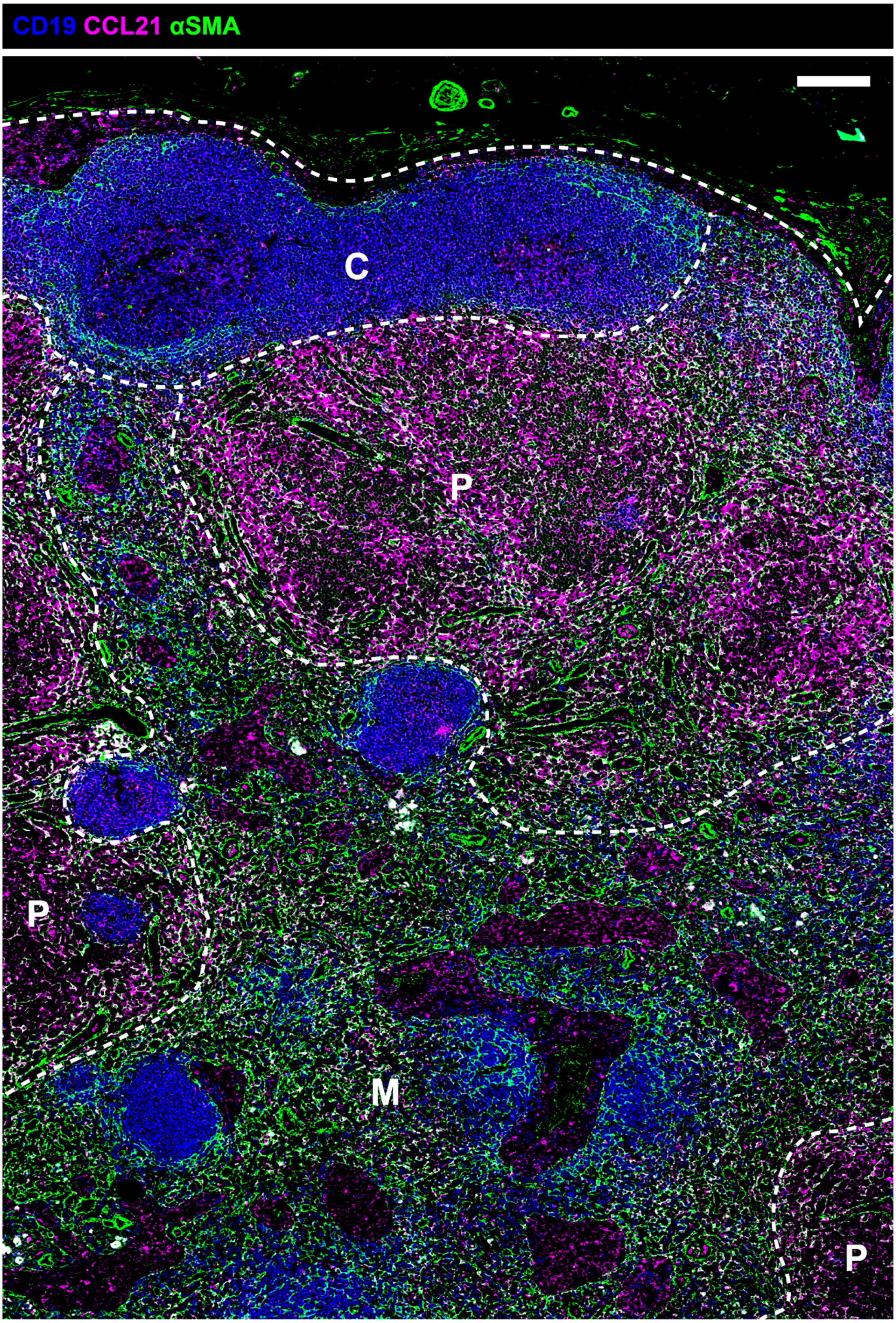


**Supplementary Figure 2: Visualization of different fibroblast subsets of the human lymph node.** Immunofluorescence staining of B-cells (CD19; blue) marking the cortex (C) and part of medulla (M), the chemokine CCL21 (magenta), marking the fibroblastic reticular cells also known as T-cell reticular cells (TRCs) of the paracortex (P), and alpha smooth muscle actin (αSMA; green) marking both the TRCs of the paracortex (P) and the medullary reticular cells (MedRC) in the medulla (M). MedRCs are low or negative for CCL21, in contrast to TRCs that express high levels. Scale bar: 150μm.

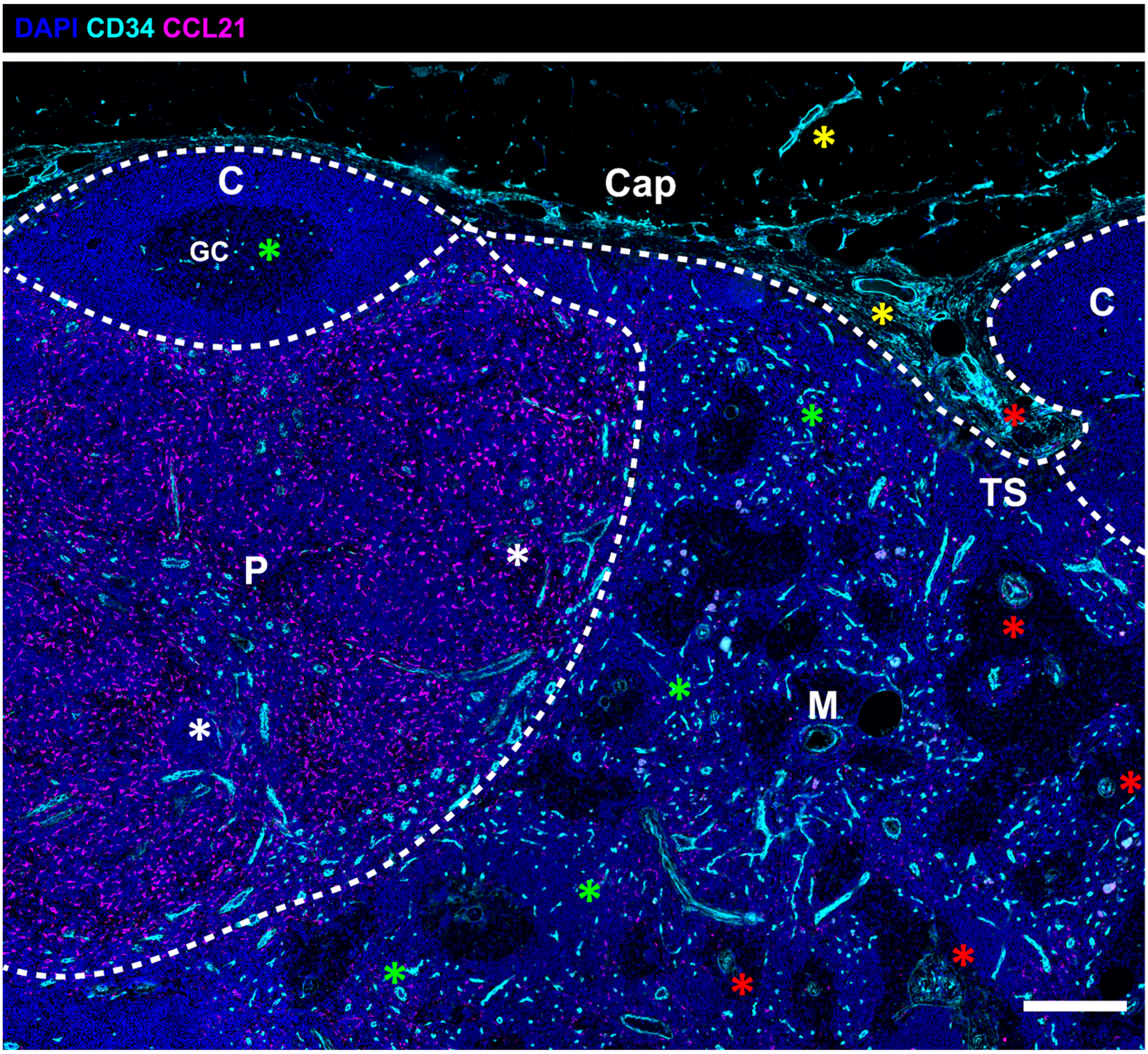


**Supplementary Figure 3: Expression pattern of CD34 in the human lymph node.** Immunofluorescent staining of a lymph node with low lipomatosis stained for CD34 (cyan), CCL21 (staining the paracortex, magenta) and DAPI (blue). CD34 is expressed in the fibroblasts of the capsule (Cap) and its invaginations called trabecula found in the trabecular sinuses (TS) (red asterisks) and in the adventitia of larger blood vessels (yellow asterisks). It is also expressed by capillary blood vessels of the germinal center (GC) of the cortex (C) and medulla (M) (green asterisks), and by the high endothelial venules (HEVs) of the paracortex (P) (white asterisks). Scale bar 250µm.

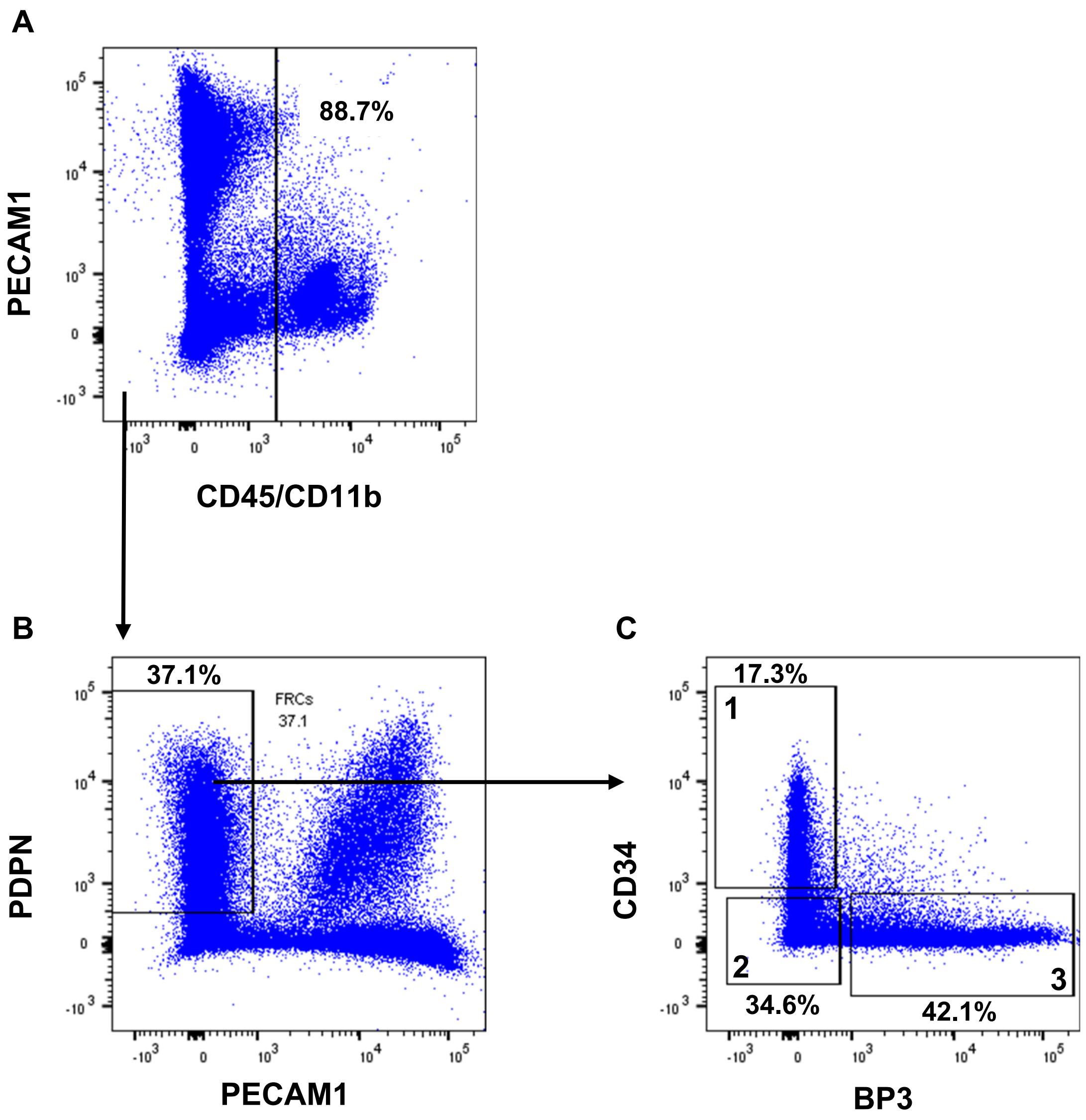


**Supplementary Figure 4: FACS**. LN fibroblasts (CD45$^{neg}$, CD11b$^{neg}$, PECAM1$^{neg}$, PDPN$^{pos.}$) were sorted into three fractions based on the markers CD34 and BP3 (Bst1): BP3$^{neg}$ CD34$^{neg}$ MedRCs, BP3$^{neg}$ CD34$^{pos}$ CD34$^{+}$ SCs and BP3$^{pos.}$ CD34$^{neg}$ RCs BP3$^{neg}$ CD34$^{neg}$ MedRCs, BP3$^{neg}$ CD34$^{pos}$ CD34$^{+}$ SCs and BP3$^{pos.}$ CD34$^{neg}$ RCs (i.e. TRCs, FDCs and MRCs). Dot plots display gating strategy and sorted populations are indicated.

A B C

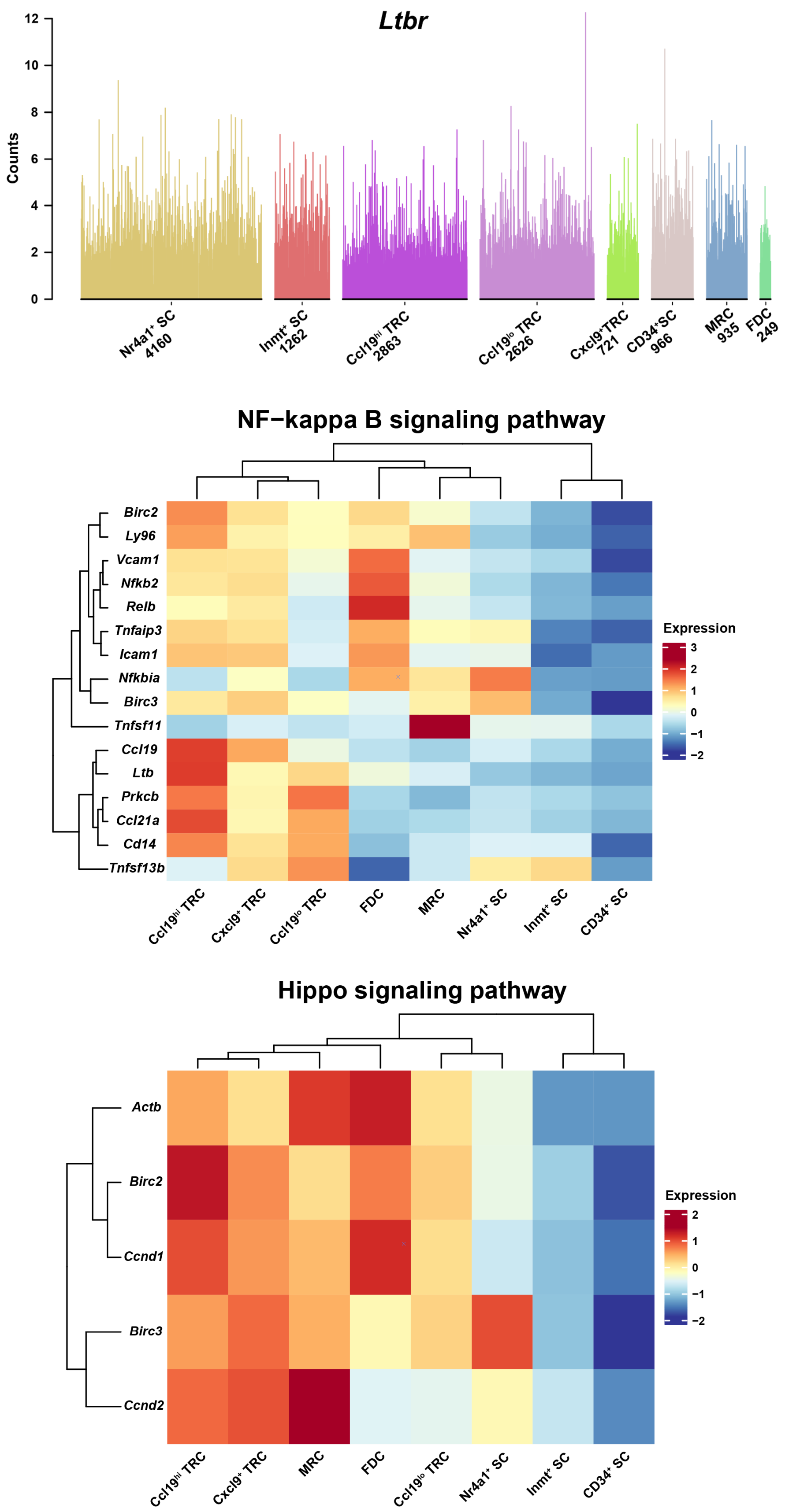


**Supplementary Figure 5: Expression analysis. (A)** Bar plot displaying expression of *Ltbr* for each cell in the different clusters described by Rodda et al., (10). (**B**) Heatmap of DEG genes ( “CD34$^{+}$ SC” cluster expressing low Bst1 vs. cell clusters expressing high Bst1) in the NFkappaB pathway, differently expressed in subsets described by Rodda et al., Values are normalized counts (log-scaled). (**C**) Heamap of DEG genes ( “CD34$^{+}$ SC” cluster expressing low Bst1 vs. cell clusters expressing high Bst1) in the Hippo pathway, differently expressed in subsets described by Rodda *et al*., Values are normalized counts (log-scaled). The MedRCs terminology, used in other parts of this paper includes the two subsets Nr4a1+ SC and Inmt+ SC, which are both BP3$^{neg/low}$ CD34$^{neg}$, the three TRC subsets, FDCs and MRCs are all BP3 high subsets, while CD34+ SC are BP3 low/neg.

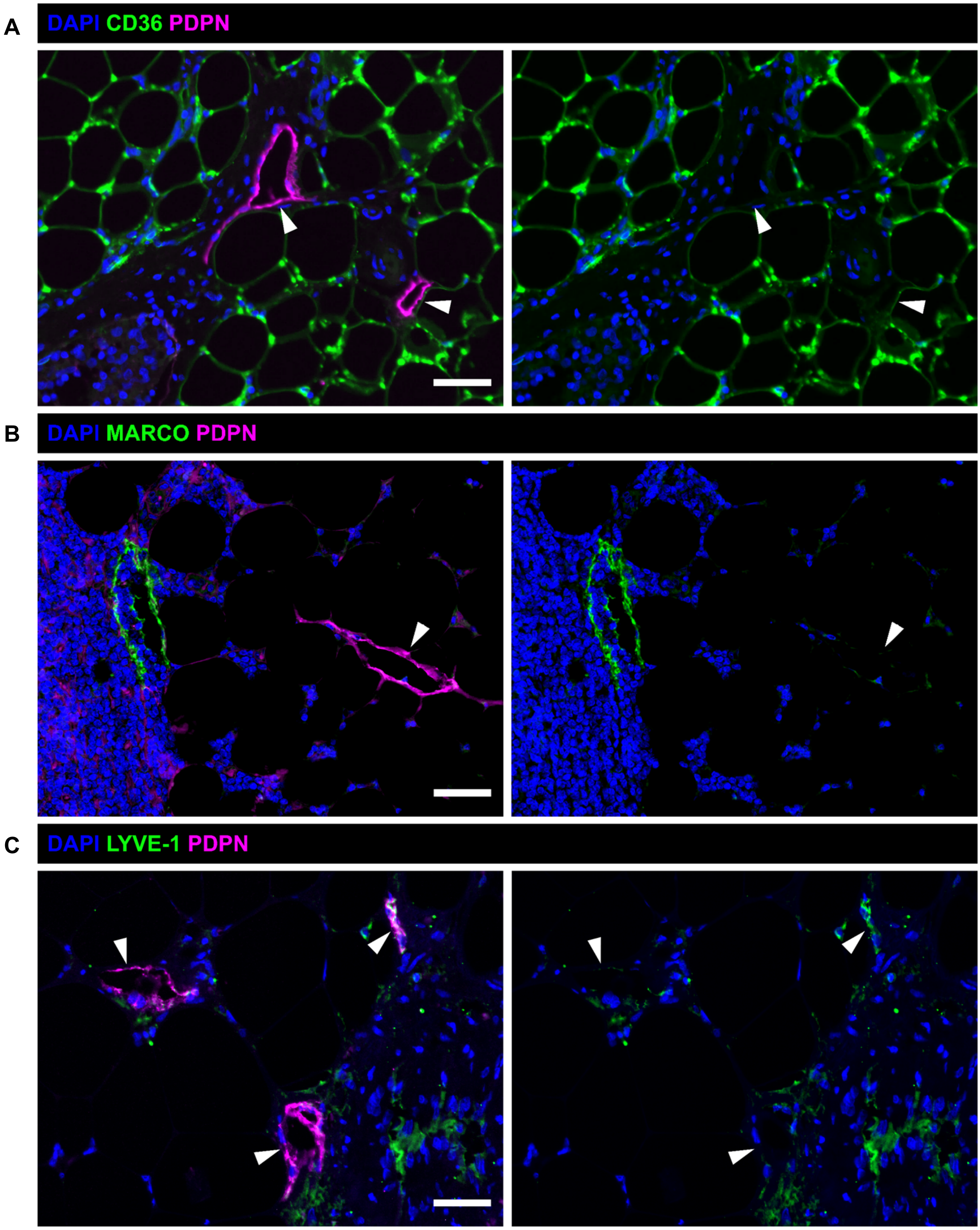


**Supplementary Figure 6. CD36, MARCO and LYVE-1 expression in collecting-like vessels.** Immunofluorescence images of lymph nodes (LNs) with intermediate to high lipomatosis. **(A)** LN stained for CD36 (green) and podoplanin (PDPN, magenta). **(B)** LN stained for MARCO (green) and PDPN (magenta). **(C)** LN stained for LYVE-1 (green) and PDPN (magenta). White arrowheads points out collecting-like vessels. Scale bar: 50µm.

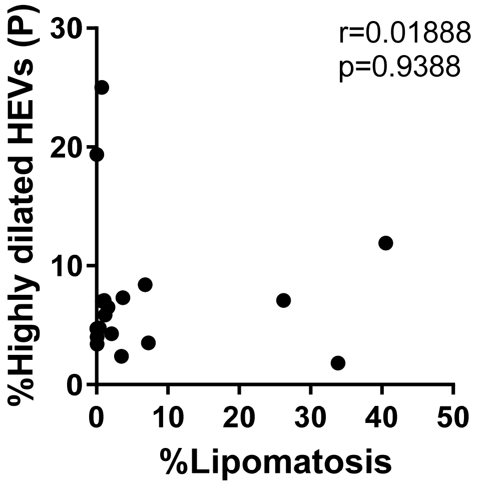


**Supplementary Figure 7: Lipomatosis is inducing a local remodeling of the high endothelial venules.** Correlation analysis of the percentage of highly dilated high endothelial venules (HEVs) in the paracortex (more than 200µm from the adipocytes) vs. the percentage of lipomatosis in the lymph node (n=19 patients). Spearman correlation test was used for statistical analysis. No correlation was detected; r=0.01888, p=0.9388.

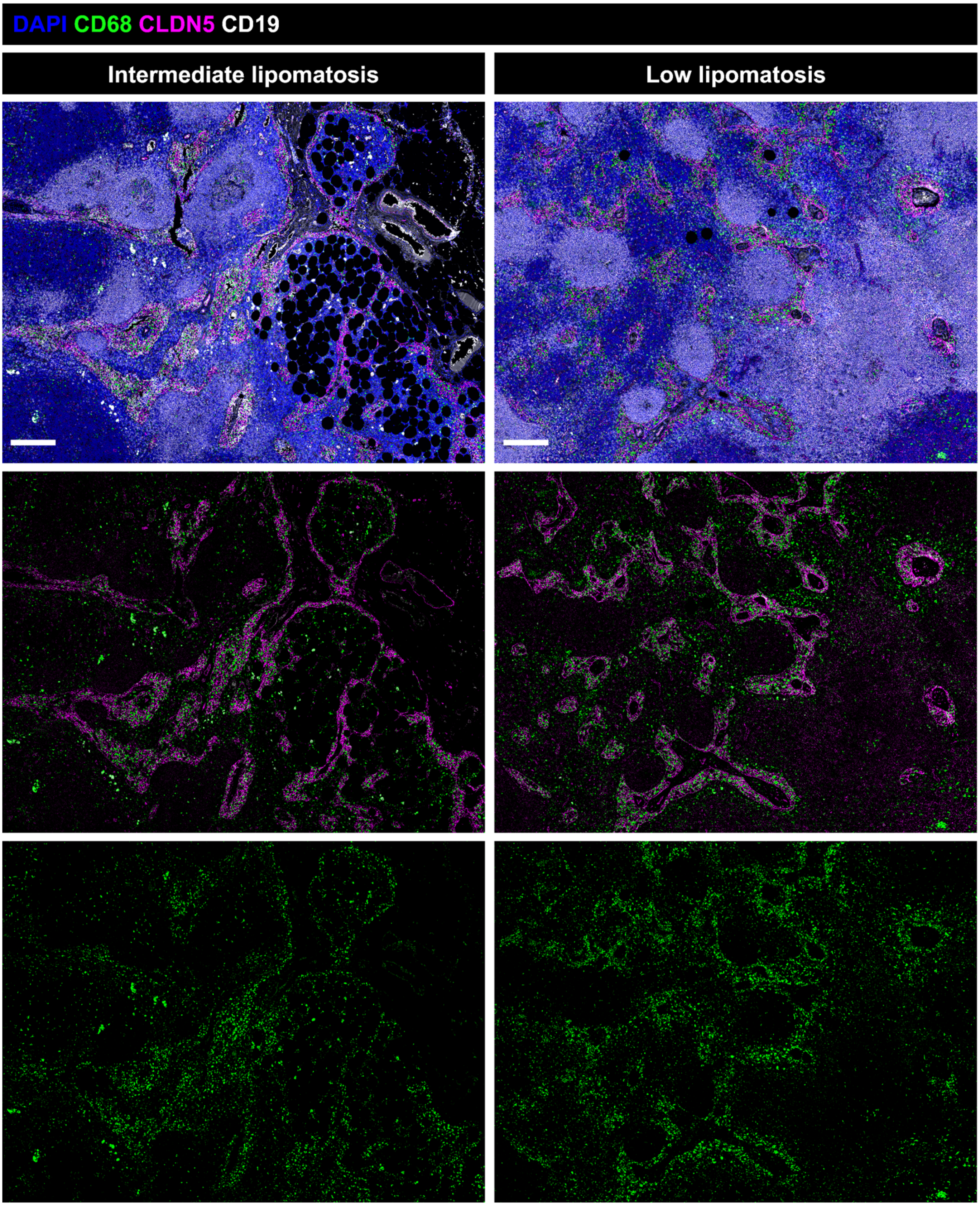


**Supplementary Figure 8: Macrophage distribution in a lipomatosis affected lymph nodes.** Immunofluorescent staining of lymph nodes with low and intermediate lipomatosis stained for the macrophage marker CD68 (green), Claudin-5 (CDLN5, magenta), CD19 (white) and DAPI (blue). Scale bar: 100µm.

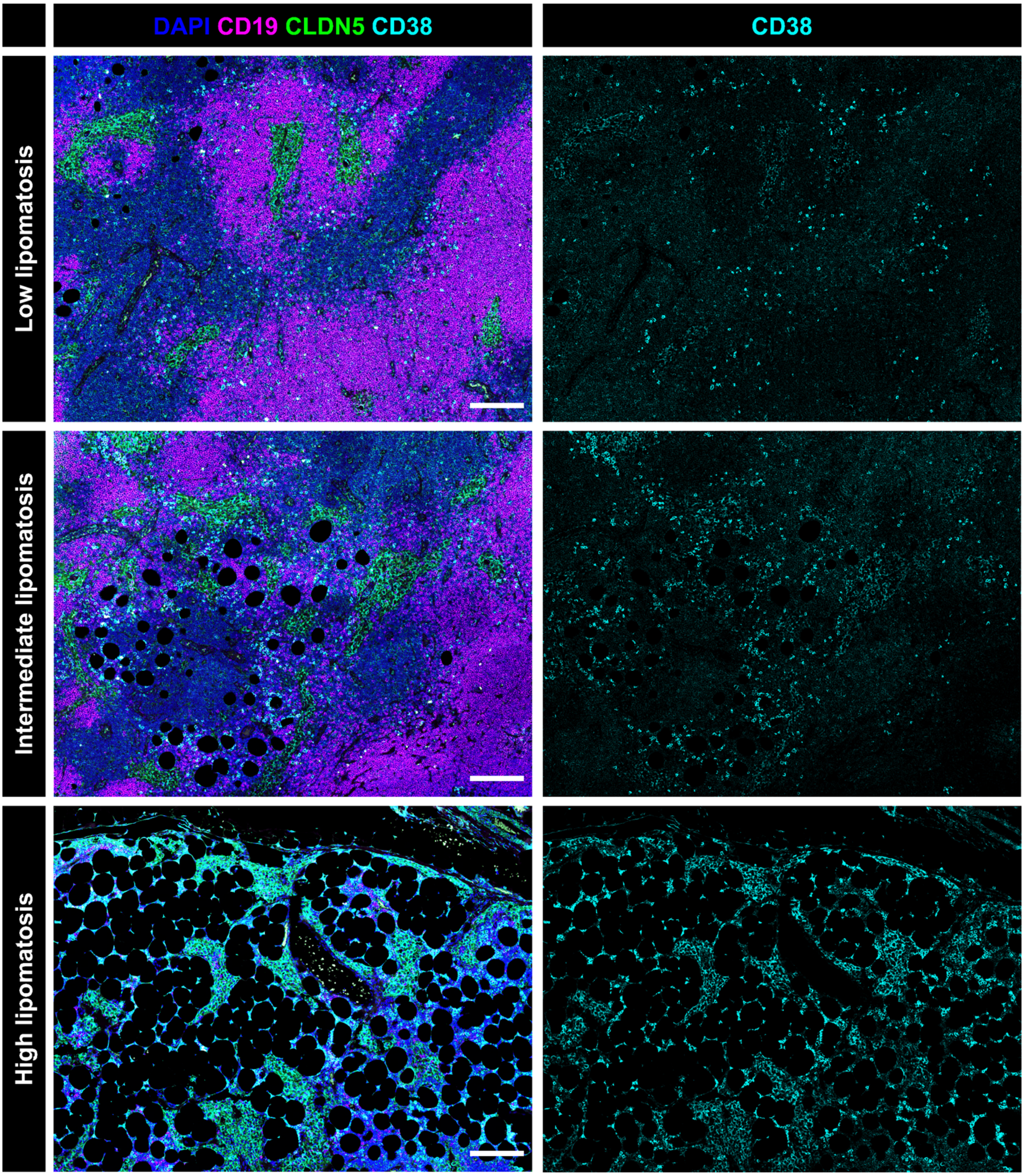


**Supplementary Figure 9: Distribution of CD38 plasma cells in the medulla of lipomatosis affected lymph nodes.** Immunofluorescent staining of CD38 (cyan), Claudin-5 (CLDN5, green), CD19 (magenta) and DAPI in areas of a lymph node (LN) with different degree of lipomatosis. Scale bar: 200µm. Pictures representative from nine LNs.
